# Topoisomerase 3B (TOP3B) DNA and RNA Cleavage Complexes and Pathway to Repair TOP3B-linked RNA and DNA Breaks

**DOI:** 10.1101/2020.03.22.002691

**Authors:** Sourav Saha, Yilun Sun, Shar-Yin Huang, Ukhyun Jo, Hongliang Zhang, Yuk-Ching Tse-Dinh, Yves Pommier

## Abstract

Genetic inactivation of TOP3B is linked with schizophrenia, autism, intellectual disability and cancer. The present study demonstrates that *in vivo* TOP3B forms both RNA and DNA cleavage complexes (TOP3Bccs) and reveals a pathway for repairing TOP3Bccs. For detecting cellular TOP3Bccs, we engineered a “self-trapping” mutant of TOP3B (R338W TOP3B) and to determine how human cells repair TOP3Bccs, we depleted tyrosyl-DNA phosphodiesterases (TDP1 and TDP2). TDP2-deficient cells produced elevated TOP3Bccs both in DNA and RNA. Conversely, overexpression of TDP2 lowered cellular TOP3Bccs. Using recombinant human TDP2, we demonstrate that TDP2 cannot excise the native form of TOP3Bccs. Hypothesizing that TDP2 cannot access phosphotyrosyl linkage unless TOP3B is either proteolyzed or denatured, we found that cellular TOP3Bccs are ubiquitinated by the E3 Ubiquitin Ligase TRIM41 before undergoing proteasomal degradation and excision by TDP2.

**HIGHLIGHTS:**

- Method for *in vivo* detection of TOP3B cleavage complexes (TOP3Bccs) formed both in DNA and RNA, using a religation defective “self-trapping” R338W TOP3B mutant.
- First evidence that TDP2 excises TOPccs produced by a type IA topoisomerase.
- TDP2 processes both RNA and DNA TOP3Bccs following their ubiquitylation and proteasomal degradation inside cell.
- TRIM41 is the first reported E3 ubiquitin ligase for TOP3Bcc ubiquitylation and proteasomal degradation.

## INTRODUCTION

Topoisomerases solve the topological constraints of nucleic acids and help during replication, transcription, recombination, chromosome segregation and chromatin remodeling. Topoisomerases act by forming transient enzyme-nucleic acid intermediate via a covalent phosphodiester bond between their catalytic tyrosine residue and one of the ends of the broken nucleic acid (3’-end for type IB and 5’-end for type IA and type II topoisomerases). These covalent catalytic intermediates are referred to as “Topoisomerase cleavage complexes” (TOPccs). Normal TOPccs are readily reversible, as they allow the resealing of the nucleic acids backbone after the topological changes and the release of topoisomerases for their next catalytic cycle (Huang and Pommier, 2019; Pommier et al., 2016; Vos et al., 2011; Wang, 2002).

When TOPccs fail to reseal, stable TOPccs are formed inside cells. DNA alterations and damages including oxidized bases, nicks, abasic sites, misincorporated ribonucleotide that occur frequently within the cell and external agents including carcinogens and anti-cancer drugs (e.g. camptothecin derivatives topotecan, irinotecan for TOP1 and doxorubicin, etoposide, amsacrine for TOP2) that binds at the topoisomerase-nucleic acid interface stabilize and enhance cellular TOPccs (Huang and Pommier, 2019; Pommier et al., 2016). These abortive TOPccs are genotoxic for cells (Pommier, 2006; Pommier et al., 2014). Eukaryotic cells deploy two alternate repair pathways to excise TOP1ccs and TOP2ccs: the tyrosyl DNA phosphodiesterase (TDP) excision pathway and the nuclease pathways (Pommier, 2006; Pommier et al., 2016). One way to repair TOP1ccs and TOP2ccs is by the action of different nucleases that cleave the DNA strand to which the topoisomerases are linked covalently (Menon and Povirk, 2016; Pommier et al., 2016). The 3’-flap endonuclease complex Rad1–Rad10 (XPF–ERCC1) has been associated with the removal of TOP1ccs (Liu et al., 2002; Vance and Wilson, 2002; Zhang et al., 2011), the 5’-flap endonuclease XPG for TOP2ccs (Kametani et al., 2016; Malik and Nitiss, 2004) and the 3’–5’ exonuclease Mre11 for both TOP1ccs and TOP2ccs (Hamilton and Maizels, 2010; Sacho and Maizels, 2011). The alternative and more specific ways to excise TOPccs involves the TDP excision pathways. TDP1 and TDP2 are two enzymes that hydrolyze the 3′-tyrosyl-DNA and 5′-tyrosyl-DNA phosphodiester bonds involving TOP1 and TOP2 respectively (Cortes Ledesma et al., 2009; Pommier et al., 2014; Pommier et al., 2016; Pouliot et al., 1999). To gain access to those phosphotyrosyl bonds, TDPs require the prior degradation or denaturation of the covalently bound topoisomerases (Pommier et al., 2016; Schellenberg et al., 2017). The ubiquitin-proteasome pathways play a pivotal role in the proteolysis of TOP1ccs and TOP2ccs in eukaryotic cells (Desai et al., 1997; Mao et al., 2001; Zhang et al., 2006) and the TDP excision pathways also depend on ubiquitin–proteasome for TOP1cc and TOP2cc repair (Pommier et al., 2016). Despite extensive research on *in vivo* detection and repair of eukaryotic type IB and type II TOPccs, little is known about eukaryotic type IA TOPccs.

Human TOP3B remained remarkably understudied until the recent discovery that it can act as a Type IA topoisomerase for both DNA and RNA (Ahmad et al., 2017a; Ahmad et al., 2017b; Ahmad et al., 2016; Stoll et al., 2013; Xu et al., 2013). Although TOP3B is not essential in flies or mice, Top3B knockout mice develop autoimmunity (Kwan et al., 2007), infertility (Kwan et al., 2003), have reduced lifespan (Kwan and Wang, 2001), abnormal neurodevelopment and defective synapse formation (Ahmad et al., 2017a; Ahmad et al., 2017b; Xu et al., 2013). In humans, TOP3B genomic deletion has been linked with Schizophrenia in a Northern Finnish subpopulation (Stoll et al., 2013). TOP3B gene deletion is also a risk factor for autism spectrum disorders (ASD) (Stoll et al., 2013; Xu et al., 2013) and TOP3B deletion has recently been observed in patients with autism, Juvenile Myoclonic Epilepsy, cognitive impairment, facial dysmorphism and behavior concerns (Ahmad et al., 2017b; Daghsni et al., 2018; Kaufman et al., 2016). In addition, two *de novo* TOP3B single nucleotide variants (SNVs), C666R and R472Q, have been reported in autism and schizophrenia patients, respectively (Iossifov et al., 2012; Xu et al., 2012). Finally, TOP3B is also linked with breast cancer (Oliveira-Costa et al., 2010) and very recently homozygous deletion for the TOP3B gene was identified in a patient with bilateral renal cancer (Zhang et al., 2019).

Little is known about the molecular mechanisms linking TOP3B to the above-mentioned diseases. TOP3B localizes to both the nucleus and cytoplasm under steady-state condition (more abundant in the cytosol) as it facilitates both DNA and RNA metabolic processes inside cells (Stoll et al., 2013; Xu et al., 2013). Inside the nucleus, TOP3B in association with the scaffolding protein TDRD3 (Tudor Domain Containing 3) is recruited to promoter regions of target genes (*NRAS, DDX5* and *c-MYC*) and has been proposed to facilitate transcription by relaxing hypernegatively supercoiled DNA and resolving R-loops (non-canonical nucleic acid structure consisting of RNA– DNA hybrids and displaced single-stranded DNA) (Goto-Ito et al., 2017; Huang et al., 2018a; Siaw et al., 2016; Yang et al., 2014; Zhang et al., 2019). TDRD3 also associates with the other well-known mRNA-binding protein FMRP (Fragile X Mental Retardation Protein) having roles in neuronal translation via its C-terminal FMRP Interacting Motif (FIM). Together the three proteins form a large heterotrimeric complex [TOP3B-TDRD3-FMRP] (Lee et al., 2018; Stoll et al., 2013; Xu et al., 2013), which remains associated with RNA during its biogenesis and maturation in eukaryotic cells (Lee et al., 2018; Stoll et al., 2013; Xu et al., 2013). Whether TOP3B catalytically engages with RNA and DNA inside cell during nucleic acid metabolic processes, is however still largely unclear and there is no direct evidence showing *in vivo* detection of catalytic intermediates of TOP3B (TOP3Bccs).

The present study introduces a novel approach to establish the presence of TOP3Bccs in human cells. It provides the first direct evidence that TOP3Bccs are formed both in cellular DNA and RNA and insights about a molecular pathway for the cellular excision of abortive TOP3Bccs. Using both biochemical and cellular assays, herein we uncover novel roles of TOP3Bcc ubiquitylation by TRIM41, proteasomal degradation of TOP3B, and of TDP2 in processing both RNA and DNA TOP3Bccs.

## RESULTS

### Generation of self-trapping mutant of TOP3B for *in vivo* detection of TOP3B cleavage complexes (TOP3Bccs)

To detect TOP3Bccs we generated a TOP3B mutant prone to remaining covalently linked to nucleic acids. Previous studies with *E. coli* and *Y. pestis* topoisomerase I (*E. coli* Topo I and *Y. pestis* Topo I) showed that substituting a single conserved arginine residues (Arg 321 in *E. coli* Topo I and Arg327 of *Y. pestis* Topo I) in the active site with a hydrophobic amino acid inhibits the resealing step of the enzyme catalytic cycle and stabilizes Topo I cleavage complexes (Narula et al., 2011). Sequence alignment (Figure 1A) of the active site regions of *E. coli* Topo I and *Y. pestis* Topo I with the corresponding region of human TOP3B revealed that Arg321 of *E. coli* Topo I and Arg327 of *Y. pestis* Topo I are conserved in human TOP3B (Arg338/R338). Hence, we hypothesized that mutating this arginine (R338) to a hydrophobic residue (e.g. tryptophan) might generate religation defective, self-trapping human TOP3B. To test this hypothesis, we generated R338W TOP3B by replacing arginine with tryptophan and transfected human HEK293 and HCT116 cells with FLAG-tagged WT and mutant TOP3B (R338W TOP3B) constructs. After 3 days of transfection, WT and R338W TOP3B proteins were readily detectable by Western blotting (Figure 1C). We next examined whether ectopic expression of R338W TOP3B induces TOP3Bccs in cells. RADAR assays (Kiianitsa and Maizels, 2013) to isolate nucleic acids with covalently-bound protein adducts showed that TOP3Bccs were readily detectable for R338W TOP3B both in HEK293 and HCT116 cells (Figures 1D and 1E).

**Figure 1.**
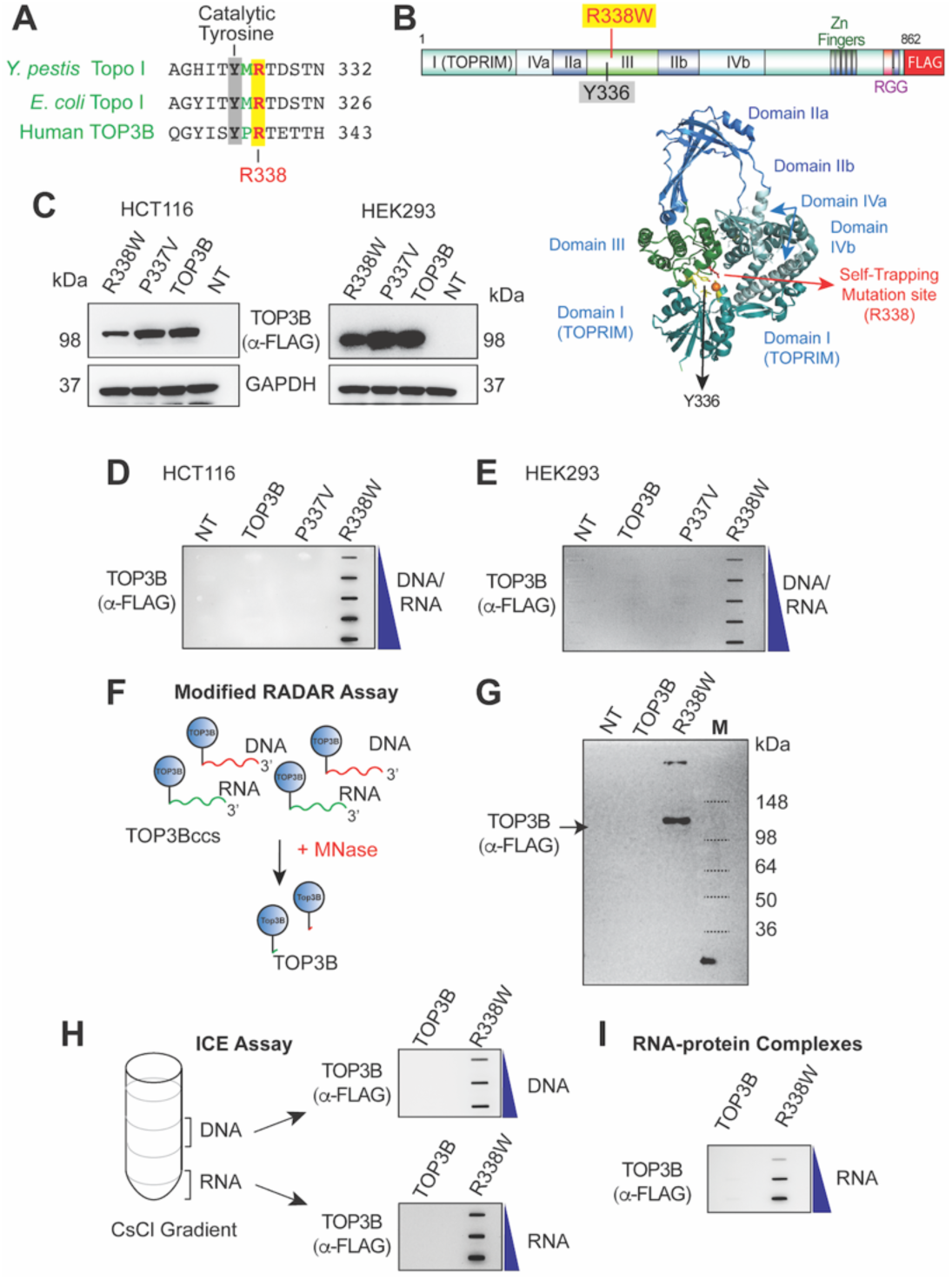
TOP3B Forms TOP3Bccs both with DNA and RNA in cells Transfected with R338W TOP3B. **(A)** Alignment of the active site regions of *E. coli* Topo I and *Y. pestis* Topo I with corresponding region of Human TOP3B. **(B)** Schematic representation of the domain organization of human TOP3B (1-862 aa) (top) and ribbon representation of human TOP3B (residues 1-612 aa) based on X-ray crystal structure by (Goto-Ito et al., 2017)(bottom). Positions of the active site residue (Tyrosine 336) and the self-trapping mutation site (Arginine 338) are indicated. **(C)** Western Blots showing ectopic expression of wild-type, P337V and R338W TOP3B. HEK293 and HCT116 cells were transfected with the indicated FLAG-tagged TOP3B constructs for 72 h and subjected to Western blotting with anti-FLAG antibody. **(D)-(E)** Detection of TOP3Bccs by RADAR assay in HEK293 and HCT116 cells transfected with FLAG-tagged wild-type, P337V or R338W TOP3B plasmid constructs for 72 h. Cells were lysed with DNAzol, nucleic acids and protein-nucleic acid adducts were isolated, slot-blotted and TOP3Bccs were detected with anti-FLAG antibody. The figure is representative of three independent experiments. **(F)** Scheme of the modified RADAR assay. Cells transfected with FLAG-tagged TOP3B (blue Circle) form TOP3Bccs in DNA (red) and RNA (green). Nucleic acids containing TOP3Bccs were isolated using the RADAR assay and digested with micrococcal nuclease (MNase) followed by SDS-PAGE and immunoblotting with anti-FLAG antibody. **(G)** Modified RADAR assay in HEK293 cells transfected with wild-type or R338W TOP3B for 72 h. TOP3B was detected with anti-FLAG antibody. **(H)** Detection of TOP3Bccs both in DNA and RNA of HEK293 cells transfected for 72 h with wild-type or R338W TOP3B constructs. Equal numbers of cells were lysed in 1% sarkosyl and ICE (In vivo complex of enzymes) bioassay by cesium chloride gradient ultracentrifugation was performed to separate DNA (middle of the gradient) and RNA (bottom of the gradient) from free proteins (top of the gradient). DNA and RNA fractions were then slot-blotted. TOP3Bccs were detected using anti-FLAG antibody. The figure is representative of three independent experiments. **(I)** Detection of RNA TOP3Bccs. HEK293 cells were transfected with wild-type or R338W TOP3B. 72 h later, covalent protein-RNA adducts were isolated using TRIzol® (Thermo Scientific), slot-blotted and TOP3Bccs were detected with anti-FLAG Antibody. The figure represents three independent experiments.

To confirm the specificity of the TOP3Bccs detected by RADAR assay, nucleic acid-containing protein adducts isolated using the RADAR assay protocol were digested with micrococcal nuclease (MNase) followed by SDS-PAGE electrophoresis and immunoblotting with anti-FLAG antibody (Figure 1F). Detection of a protein band corresponding to the size of human TOP3B (Figure 1G) indicated that R338W TOP3B formed TOP3Bccs inside cells.

### TOP3B forms both DNA and RNA cleavage complexes (TOP3Bccs) *in vivo*

Next, to determine whether TOP3Bccs form both on DNA and RNA inside cells, we transfected HEK293 cells with FLAG-tagged TOP3B and R338W TOP3B constructs and performed ICE (In vivo complex of enzymes) bioassays (Figure 1H). DNA- and RNA-protein adducts were separated from the free proteins by cesium chloride gradient ultracentrifugation and slot-blots of isolated DNA- and RNA-protein adducts were performed. TOP3Bccs were present both in the DNA and RNA fractions (Figure 1H).

To confirm this finding, we ectopically expressed R338W TOP3B in HEK293 cells and isolated nucleic acid-containing protein adducts by RADAR assay. Equal amounts of RADAR assay samples were digested either with excess RNase A and RNase T1 to remove RNA, or with DNase I to digest DNA away or with micrococcal nuclease (MNase) to remove total nucleic acids (Supplemental Figure 1A). Samples were slot-blotted and TOP3Bccs detected using anti-FLAG antibody. TOP3Bccs were readily detectable after digestion with RNase A and RNase T1 mix or DNase I (Supplemental Figure 1A and 1B), which substantiates our finding that TOP3B forms both DNA and RNA TOP3Bccs inside cells.

To further establish the formation of TOP3Bccs on RNA, we isolated covalent RNA-protein adducts from cells transfected with R338W TOP3B construct using TRIzol® reagent. Slot blotting of isolated RNA-protein adducts confirmed the formation of RNA-TOP3Bccs *in vivo* (Figure 1I).

### TDP2 excises both DNA and RNA TOP3Bccs) in human cells

In eukaryotes, irreversible TOPccs are processed by TDP1 and TDP2. To determine whether these tyrosyl phosphodiesterases repair TOP3Bccs, first we individually knocked down TDP1 and TDP2 or both TDP1 and TDP2 together using siRNAs (Supplemental Figures 2A and 2B) in HEK293 cells transfected with FLAG-tagged R338W TOP3B. RADAR assays showed that TDP2 knockdown and dual TDP1 and TDP2 knockdown cells displayed significant increase in TOP3Bccs whereas TDP1 knockdown cells accumulated similar level of TOP3Bcc compared to control cells (Figure 2A and 2B). This indicated a role for TDP2 in processing of TOP3Bccs.

**Figure 2.**
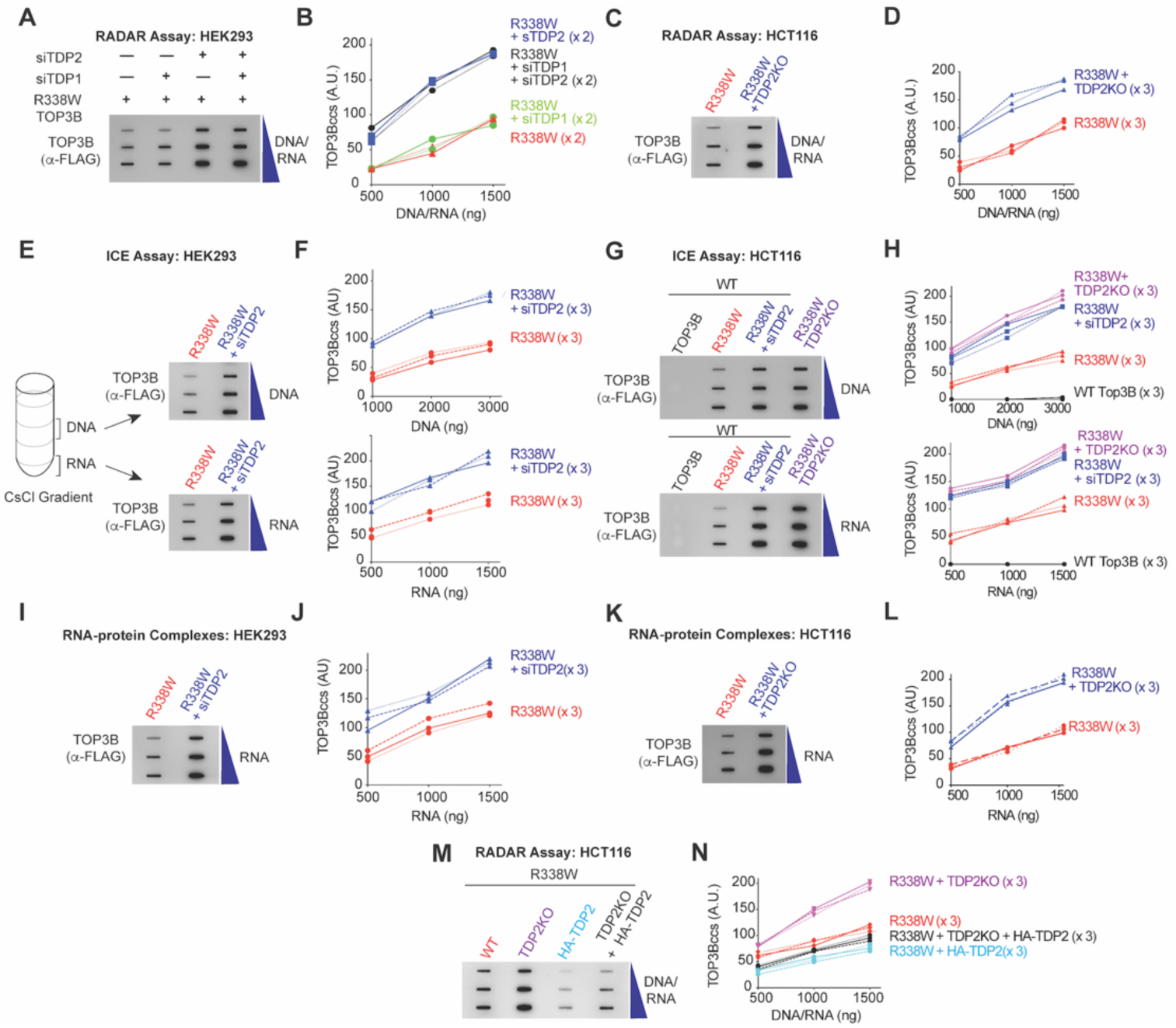
TDP2 Excises Cellular TOP3Bccs both from DNA and RNA. **(A-D)** DNA & RNA TOP3Bccs detection by RADAR assays. (A) HEK293 cells were transfected with the indicated siRNA: siTDP1, siTDP2 or both siTDP1 and siTDP2 and co-transfected with R338W TOP3B. After 72 h, nucleic acids and protein-nucleic acid adducts were isolated by RADAR assay, slot-blotted and detected with anti-FLAG antibody. The figure is representative of two independent experiments. (B) Quantitation of TOP3Bcc of RADAR assays as shown in panel A. TOP3Bccs were measured by densitometric analyses of slot-blot signals and plotted as a function of total nucleic acid (DNA and RNA) concentration. Two independent experiments are plotted. (C) Wild-type or TDP2KO HCT116 cells were transfected with FLAG-tagged R338W TOP3B. After 72 h, nucleic acids and protein-nucleic acid adducts were isolated by RADAR assay and slot-blotted. TOP3Bccs were detected using anti-FLAG antibody. The figure is representative of three independent experiments. (D) Quantitation of TOP3Bcc formation in RADAR assays as shown in panel C. Three independent experiments are plotted. **(E-H)** DNA & RNA TOP3Bccs detection by ICE assays. (E) HEK293 cells were transfected with R338W TOP3B alone or co-transfected with siTDP2. After 72 h, ICE bioassay was performed to separate DNA and RNA from free proteins. DNA and RNA fractions were slot-blotted and TOP3Bccs were detected using anti-FLAG antibody. The figure is representative of three independent experiments. (F) Quantitation of TOP3Bcc in the DNA and RNA fractions as shown in panel E. Three independent experiments are plotted. (G) HCT116 WT and TDP2KO cells were transfected either with wild-type TOP3B or R338W TOP3B or a combination of R338W TOP3B and siTDP2 constructs as indicated. After 72 h, ICE bioassays were performed to separate DNA and RNA from free proteins. DNA and RNA fractions were slot-blotted and TOP3Bccs were detected using anti-FLAG antibody. The figure is representative of three independent experiments. (H) Quantitation of TOP3Bcc formation in ICE assays as shown in panel G. Three independent experiments are plotted. **(I-L)** Detection of RNA TOP3Bccs. (I) HEK293 cells were transfected with R338W TOP3B alone or co-transfected with siTDP2. After 72 h, covalent protein-RNA adducts were isolated using TRIzol^®^, slot-blotted and TOP3Bccs were detected using anti-FLAG Antibody. The figure is representative of three independent experiments. (J) Quantitation of TOP3Bcc in RNA as shown in panel I. Three independent experiments are plotted. (K) Wild-type and TDP2KO HCT116 cells were transfected with FLAG-tagged R338W TOP3B. After 72 h, covalent protein-RNA adducts were isolated using TRIzol^®^, slot-blotted and TOP3Bccs were detected using anti-FLAG Antibody. The figure is representative of three independent experiments. (L) Quantitation of RNA TOP3Bcc in RNA as shown in panel K. Three independent experiments are plotted. **(M-N)** Ectopic expression of TDP2 reduces TOP3Bccs. (M) Wild-type (WT) and TDP2KO HCT116 cells were transfected with FLAG-tagged R338W TOP3B alone or co-transfected with HA-tagged TDP2. After 72 h, nucleic acids and protein-nucleic acid adducts were isolated by RADAR assay, slot-blotted and TOP3Bccs were detected with anti-FLAG antibody. The figure is representative of three independent experiments. (N) Quantitation of TOP3Bcc formation using the RADAR assays as shown in panel M. Three independent experiments are plotted.

To consolidate these results, we repeated the experiments in isogenic TDP2 knockout (TDP2KO) HCT116 cells (Supplemental Figure 2C) [generated by CRISPR-Cas9 (Huang et al., 2018b)]. RADAR assays showed elevated TOP3Bccs in TDP2KO cells transfected with R338W TOP3B (Figure 2C and 2D). Together these data indicate that TDP2 eliminates TOP3Bccs inside cells.

Next, to determine whether TDP2 can excise cellular TOP3Bcc from both DNA and RNA, we performed ICE (In vivo complex of enzymes) assays to separate and examine the DNA and RNA TOP3Bcc. Knocking-down TDP2 using siRNA in HEK293 cells significantly enhanced TOP3Bcc levels both in the DNA and RNA fractions (Figure 2E and 2F). Similar results were obtained by performing ICE assays in the isogenic HCT116 TDP2KO cells (Figure 2G and 2H). To independently validate our finding that TDP2 excises TOP3Bccs both from DNA and RNA, we took equal amount of RADAR assay samples prepared from control and siTDP2-transfected HEK293 cells transiently expressing R338W TOP3B and digested them either with excess RNase A and RNase T1, or with DNase I to distinguish the DNA and RNA TOP3Bccs. Downregulation of TDP2 increased both DNA and RNA TOP3Bccs (Supplemental Figures 2D and 2E).

To further establish that depletion of TDP2 selectively enhances RNA TOP3Bccs, we isolated covalent RNA-protein adducts with TRIzol® from TDP2-knocked-down and control HEK293 cells transfected with R338W TOP3B. Downregulation of TDP2 led to a significantly increase of RNA TOP3Bccs in RNA (Figure 2I and 2J). Similarly, knocking-out TDP2 in HCT116 cells significantly enhanced TOP3Bccs in cellular RNA (Figure 2K and 2L).

We also performed TDP2 complementation experiments by ectopic expression of HA-tagged TDP2 (Supplemental Figure 2F). RADAR assays showed significantly reduced TOP3Bccs both in wild-type and TDP2KO HCT116 cells ectopically expressing HA-tagged TDP2 (Figure 2M and 2N). Together these results lead us to conclude that TDP2 has a definitive role in the excision of both cellular DNA and RNA TOP3Bccs.

### Human TDP2 cannot process Native TOP3Bccs

To determine whether recombinant TDP2 can directly hydrolyze the TOP3B phosphodiester bond and excise TOP3B from TOP3Bccs, we engineered a suicidal hairpin substrate with a long 3’-tail, which we incubated with recombinant human TOP3B to generate irreversible (suicidal) TOP3Bccs (Figure 3A). TOP3Bccs resulted in reduced mobility of TOP3B in SDS-PAGE gel (Figure 3A, lane 2). As expected, benzonase, which was used as a positive control, degraded the nucleotide and released free TOP3B (Figure 3A, lanes 3-4). However, neither TDP1 nor TDP2 could process the suicidal TOP3Bccs even at high concentration (Figure 3A, lanes 5-8). From these experiments, we conclude that TDP2 cannot excise intact, native TOP3B from TOP3Bccs.

**Figure 3.**
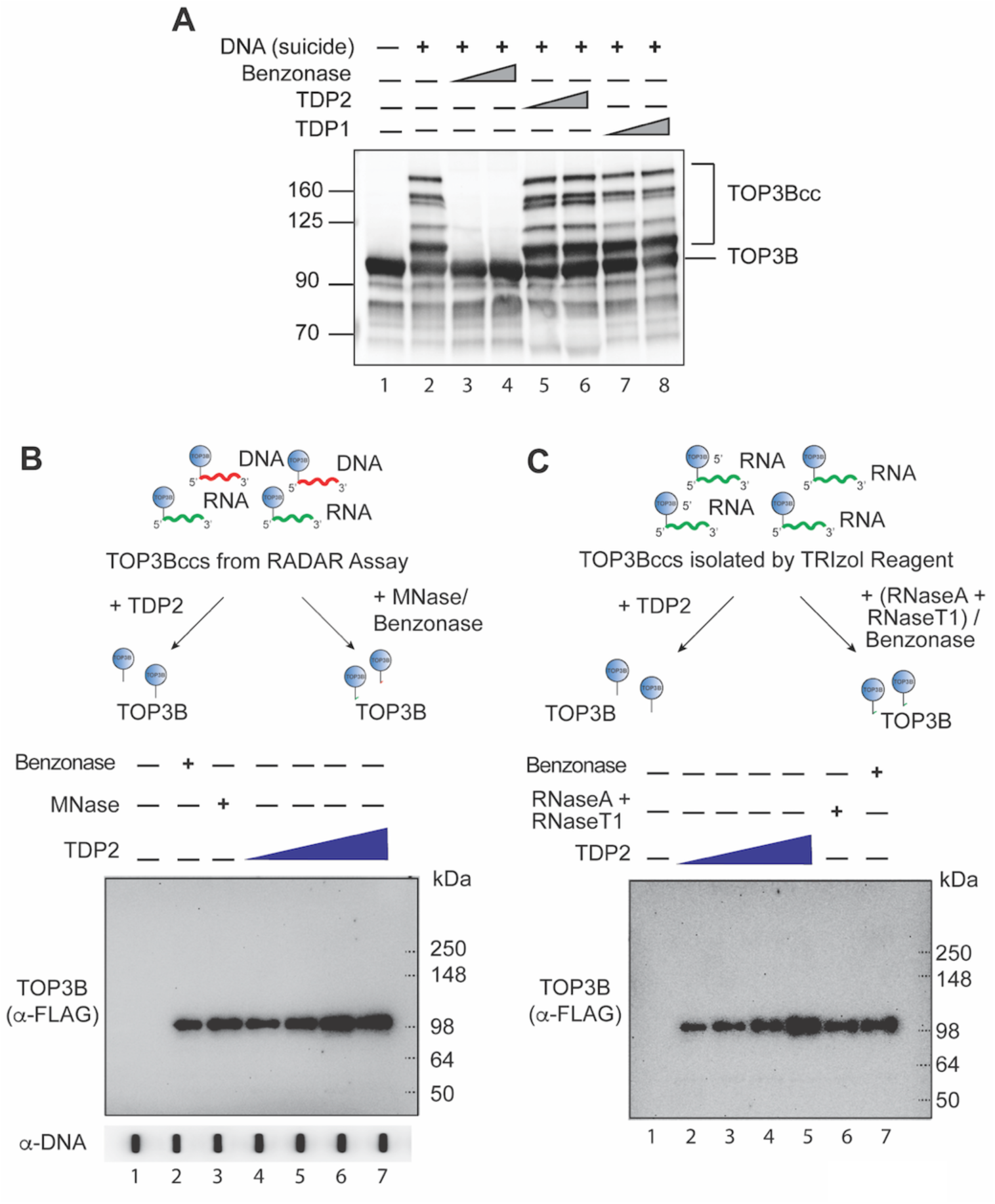
Recombinant Human TDP2 Only Unhooks Denatured but Not Native TOP3B from TOP3Bcc. **(A)** Recombinant human TDP2 does not excise native TOP3Bccs. A 69-mer single-stranded DNA oligonucleotide was designed to study the processing of suicidal TOP3Bccs by TDP2. The oligonucleotide (300 nM)was incubated with purified recombinant human TOP3B (4 uM). Irreversible (suicidal) TOP3Bccs result in slower migrating DNA (Lane 2). Positions of free TOP3B and TOP3Bccs are indicated. Suicidal TOP3Bccs were incubated with increasing concentration (1 or 3 μM) of recombinant TDP1 (Lanes 7 and 8) or TDP2 (1 or 3 μM, Lanes 5 and 6). Benzonase (3 or 9 Units, Lanes 3 and 4) was used as positive control for degradation of the oligonucleotide and release of TOP3B. Samples were resolved by SDS-PAGE and immunoblotted with anti-TOP3B antibody. **(B-C)** Recombinant human TDP2 excises denatured cellular DNA and RNA TOP3Bccs. (B) HEK293 cells were transfected with FLAG-tagged R338W TOP3B and nucleic acids and protein-nucleic acid adducts were recovered by RADAR assay. After incubation with increasing concentrations of recombinant TDP2 (1, 2, 3 and 6 μM, lanes 4-7), micrococcal nuclease (MNase, 300 Units/reaction, lane 3) or benzonase (250 Units/reaction, lane 2), reaction mixtures were analyzed by immunoblotting with anti-FLAG antibody after SDS-PAGE. Benzonase and MNase were used as controls for complete degradation of DNA and RNA in TOP3Bccs. (C) HEK293 cells were transfected with R338W TOP3B. Covalent protein-RNA adducts were isolated 72 hs later using TRIzol® reagent. After incubation with increasing concentrations of recombinant TDP2 (1, 2, 3 and 6 μM, lanes 2-5), excess amount of RNase A (200 μg/mL) and RNase T1 (200 Units/ml) mix (Lane 6) or benzonase (250 Units/ reaction, lane 7), reaction mixtures were analyzed by immunoblotting with anti-FLAG antibody after SDS-PAGE. Benzonase and RNase A and RNase T1 mix were used as controls for complete degradation of RNA covalently attached to TOP3Bccs.

We also examined whether recombinant human TDP2 could process TOP3Bccs when TOP3B is denatured. To do so, we isolated TOP3Bccs by RADAR assay from FLAG-tagged R338W TOP3B-transfected HEK293 cells. Indeed, the RADAR assay uses a combination of chaotropic salt and detergents that denature TOP3B in the TOP3Bccs. These RADAR assay samples were incubated with increasing concentration of recombinant TDP2, MNase (or benzonase and reaction mixtures were analyzed by Western blotting with anti-FLAG antibody to detect released TOP3B. As expected, untreated TOP3Bccs were unable to enter the gels due to their covalent linkage to nucleic acids (Figure 3B, lane 1). Benzonase and MNase were used as controls for complete degradation of the nucleic acids (DNA and RNA) and to release TOP3B from TOP3Bccs. After treatment with TDP2, MNase or benzonase, samples migrated as a single band (Figure 3B, lanes 2-7) corresponding to the size of FLAG-tagged TOP3B (∼100 kDa). These findings imply that recombinant TDP2 can resolve TOP3Bccs when TOP3B is denatured.

Next we tested whether the processing of denatured TOP3Bcc is also observed for RNA TOP3Bccs. We isolated the RNA TOP3Bccs from cells transiently expressing R338W TOP3B using the TRIzol® procedure. TDP2, like benzonase or RNase A and RNase T1 mix, was able to release TOP3B from denatured RNA TOP3Bccs (Figure 3C). Together, these results demonstrate that TDP2 can remove denatured but not native TOP3B from both DNA and RNA TOP3Bccs.

### Ubiquitination and proteasomal degradation of cellular TOP3Bccs

From the above results, we hypothesized that, unless TOP3B loses its native structure and/ or gets proteolyzed, TDP2 cannot access and hydrolyze the phosphotyrosyl bond between the nucleic acid and TOP3B. To determine whether ubiquitin-mediated proteasomal degradation plays a role in the processing of TOP3Bccs, we treated HEK293 and HCT116 cells transfected with FLAG-tagged R338W or wild-type TOP3B constructs with the proteasome inhibitor MG132. RADAR assays showed that MG132 increased TOP3Bccs both in the R338W TOP3B-transfected HEK293 and HCT116 cells (Figures 4A-D), demonstrating an implication of proteasomal degradation in the repair of cellular TOP3Bccs. To further establish the role of ubiquitin in TOP3Bcc repair, we treated FLAG-tagged R338W TOP3B transfected HEK293 and HCT116 cells with the E1 ubiquitin-activating enzyme (UAE) inhibitor TAK-243. RADAR assays showed that TAK-243 enhanced the cellular levels of TOP3Bccs (Figures 4E-H). These results demonstrate that ubiquitination of TOP3B is associated with the proteasomal processing of TOP3Bccs.

**Figure 4.**
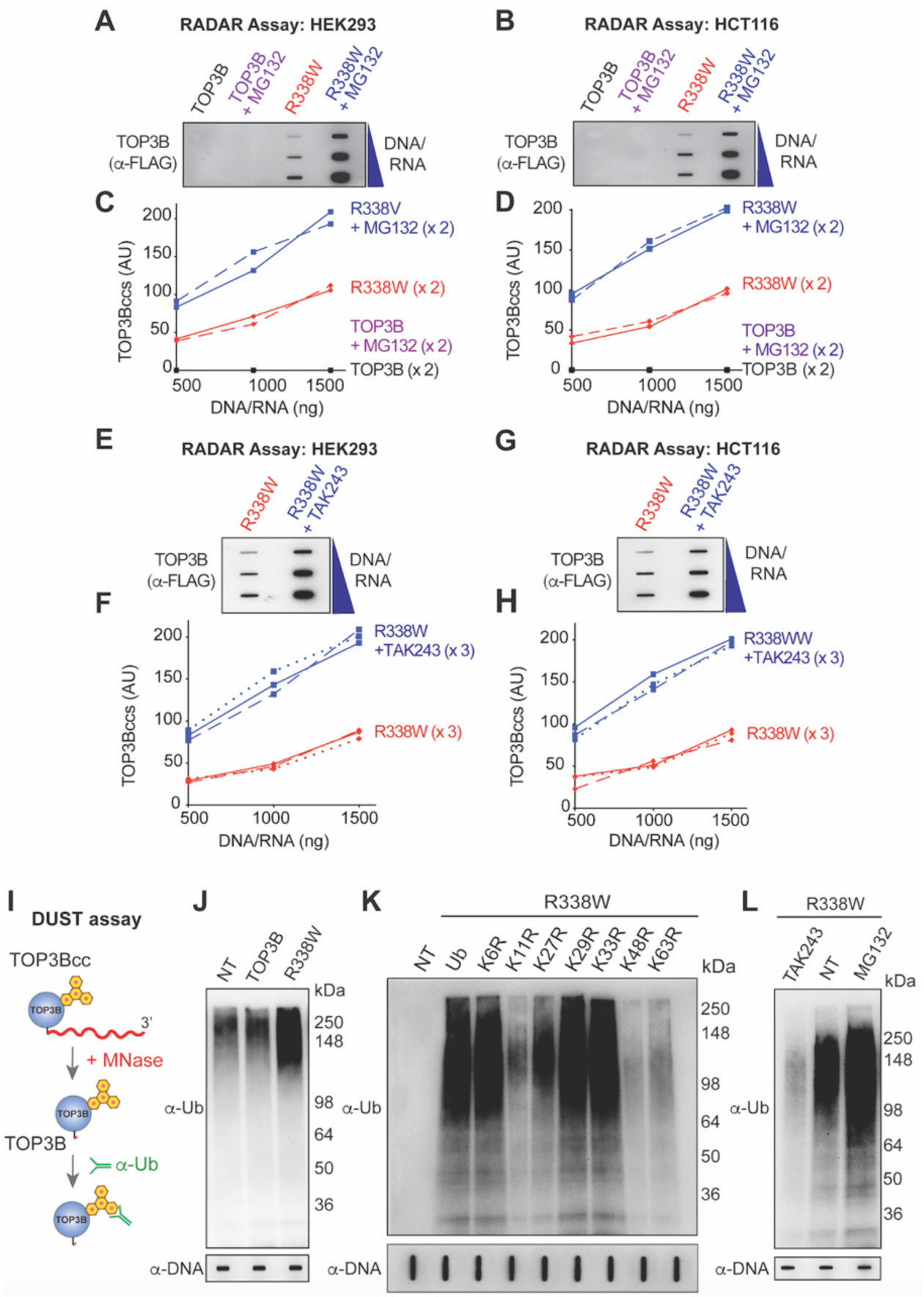
Cellular TOP3Bccs are Ubiquitylated and Degraded by the Proteasomal Pathway. **(A-D)** Proteasome inhibition enhances cellular TOP3Bccs. HEK293 (A) and HCT116 cells (B) were transfected with FLAG-tagged wild-type TOP3B and R338W TOP3B for 72 h. Before harvest, cells were treated with MG132 (10 µM, 2 h). TOP3Bccs were detected by RADAR assays using anti-FLAG antibody. The figure is representative of two independent experiments. (C-D) Quantitation of TOP3Bccs in two independent RADAR assays as shown in panels A and B. Two independent experiments are plotted for each panels. **(E-H)** Ubiquitylation inhibition enhances cellular TOP3Bccs. HEK293 (E) and HCT116 cells (G) were transfected with FLAG-tagged R338W TOP3B for 72 h. Before harvesting, the cells were treated with the UAE inhibitor TAK-243 (10 µM, 2 h). TOP3Bccs were detected by RADAR assays using anti-FLAG antibody. The figure is representative of three independent experiments. (F & H) Quantitation of RADAR assays as shown in panels E and G. Three independent experiments are plotted for each panels. **(I)** Scheme of the DUST (Detection of Ubiquitylated-SUMOylated TOPccs) assay. Following the isolation of TOP3Bccs by RADAR assay, the covalently attached nucleic acids are digested with micrococcal nuclease (MNase). Ubiquitylated TOP3B can then be detected by immunoblotting following SDS-PAGE. **(J)** Ubiquitylation of TOP3Bccs in HCT116 cells transfected with FLAG-tagged R338W TOP3B for 72 h, as detected by the DUST Assay. Equal loading was tested by slot-blotting and probing with anti-dsDNA antibody. **(K)** TOP3Bcc ubiquitylation involves the classical proteasomal-specific linkages to lysines K11, K27, K48 and K63. HCT116 cells were co-transfected with FLAG-tagged R338W TOP3B plasmid construct and HA-tagged wild-type or mutant ubiquitin constructs. DUST assays were performed after 72 h. **(L)** Inhibition of TOP3Bcc ubiquitylation by the UBE1 inhibitor TAK-243 and enhancement by the proteasome inhibitor MG132. HCT116 cells transfected with FLAG-tagged R338W TOP3B for 72 h were treated with either MG132 (10 µM, 2 h) or TAK-243 (10 µM, 2 h), as detected by the DUST Assay.

To determine whether the TOP3Bccs are ubiquitylated, we performed Detection of Ubiquitylated and SUMOylated TOP-DPCs [DUST assay (Sun et al., 2019)]. In this assay RADAR samples are digested with micrococcal nuclease (MNase) to release TOP3B from TOP3Bccs and ubiquitylated TOP3B are detected by SDS-PAGE and immunoblotting with anti-Ub antibody (Figure 4I). HCT116 cells transfected with FLAG-tagged R338W TOP3B showed enhanced cellular ubiquitylation of TOP3Bccs (compared to wild-type TOP3B transfected cells) (Figure 4J).

Ubiquitin has seven lysine residues (K6, K11, K27, K29, K33, K48 and K63), which are employed during polyubiquitin chain formation on substrate proteins. To determine the polyubiquitin linkages of TOP3Bccs, we co-transfected HCT116 cells with R338W TOP3B and either wild type HA-tagged Ub or HA-tagged lysine-to-arginine Ub mutants for each of these seven lysine residues (K6R-Ub, K11R-Ub, K27R-Ub, K29R-Ub, K33R-Ub, K48R-Ub and K63R-Ub). DUST assays revealed that K11, K27, K48 and K63 determine TOP3Bcc ubiquitylation (Figure 4K), indicating the presence of K11, K27, K48 and K63 linkages in the polyubiquitin chains associated with TOP3Bccs.

To further establish the ubiquitination and proteasomal processing of TOP3Bccs, HCT116 cells transfected with FLAG-tagged R338W TOP3B were treated with either MG132 or TAK-243 before harvest. DUST assays were performed to compare the levels of ubiquitinated TOP3Bccs under these different conditions. While proteasome inhibition by MG132 further increased ubiquitylated TOP3Bccs, TAK-243 drastically reduced ubiquitylation of TOP3Bccs (Figure 4L). Together our findings point towards the fact that the repair of trapped cellular TOP3Bccs is associated with ubiquitylation and proteasomal degradation.

### TOP3Bccs are ubiquitylated by E3 ubiquitin ligase TRIM41 in Human Cells

Tripartite motif (TRIM) proteins constitute a large class of “single protein RING finger E3 ubiquitin ligases” (Ikeda and Inoue, 2012). Because TRIM41 protein had been identified as an interaction partner of TOP3B by yeast 2-hybrid analysis (Kobayashi and Hanai, 2001) and by high throughput proteome analyses (Rolland et al., 2014), we tested whether TRIM41 could act as E3 for TOP3Bccs. Downregulation of TRIM41 by siRNA increased the total cellular level of TOP3B protein (Figure 5A) and RADAR assays showed that cells transfected with siTRIM41 accumulated more TOP3Bccs (Figures 5B and C). DUST assays also showed that TRIM41 downregulation reduced the level of ubiquitylated TOP3Bccs inside cells (Figure5D). These results indicate that TRIM41 induces the ubiquitylation, degradation and excision of TOP3Bccs.

**Figure 5.**
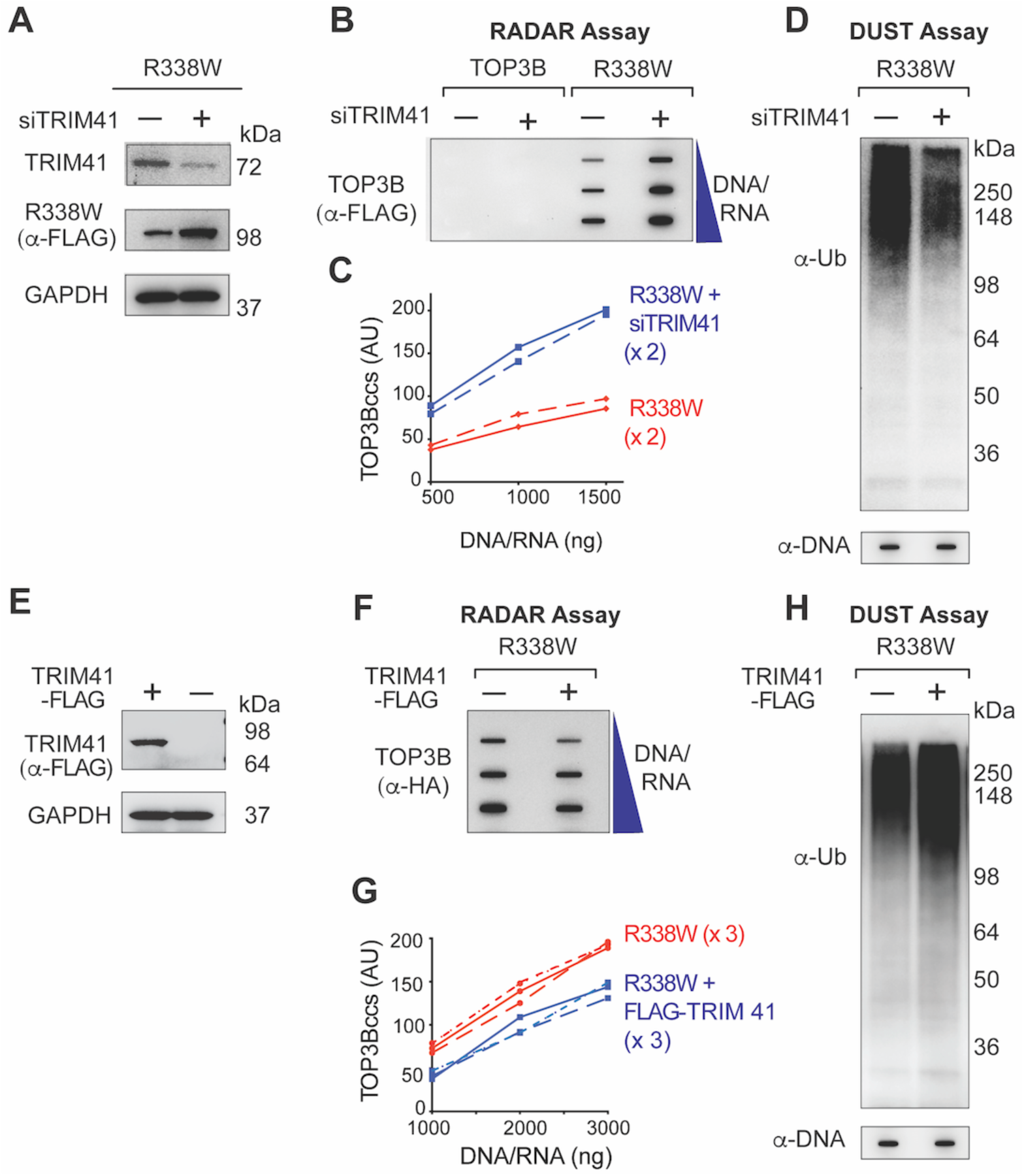
TRIM41 acts as Ubiquitin Ligase for TOP3Bccs and Promotes the Repair of TOP3Bccs. **(A)** Immunoblots showing TRIM41 and R338W TOP3B expression after TRIM41 down-regulation (GAPDH as loading control). HCT116 cells were either transfected with FLAG-tagged R338W TOP3B plasmid construct alone or co-transfected with siTRIM41construct for 72 h. **(B)** HCT116 cells were transfected with FLAG-tagged wild-type or R338W TOP3Bs or co-transfected with siTRIM41construct for 72 h. Cells were harvested and nucleic acids and protein-nucleic acid adducts isolated by RADAR assay. TOP3Bcc were detected with anti-FLAG antibody. The figure is representative of two independent experiments. **(C)** Quantitation of TOP3Bccs in 2 independent RADAR assays as shown in panel B. **(D)** HCT116 cells were transfected with FLAG-tagged R338W TOP3B alone or with siTRIM41construct. After 72 h, TOP3Bcc ubiquitylation was detected by DUST assay. Samples were also subjected to slot-blotting and probed with anti-dsDNA antibody as loading control. **(E)** Immunoblot showing expression of TRIM41 with GAPDH as loading control. HCT116 cells transfected with FLAG-tagged TRIM41 for 48 h. **(F)** HCT116 cells were either transfected with HA-tagged R338W TOP3B plasmid construct alone or co-transfected with FLAG-tagged TRIM41 construct. After 48 h cells were harvested and nucleic acids and protein-nucleic acid adducts were isolated by RADAR assay. TOP3Bccs were detected using anti-HA antibody. The figure is representative of three independent experiments. **(G)** Quantitation of TOP3Bcc formation in 3 independent RADAR assays as shown in panel F. **(H)** HCT116 cells were either transfected with HA-tagged R338W TOP3B plasmid construct alone or co-transfected with FLAG-tagged TRIM41construct. After 48 h, TOP3Bcc ubiquitylation was detected by DUST assay. Samples were also subjected to slot-blotting and probing with anti-dsDNA antibody as loading control.

To further validate the implication of TRIM41 in the repair of TOP3Bccs, we overexpressed FLAG-tagged TRIM41 in HCT116 cells transiently harboring HA-tagged R338W TOP3B. Ectopic expression of TRIM41 was confirmed by Western blot using anti-FLAG antibody (Figure 5E) and RADAR assays showed that overexpression of TRIM41 decreased cellular TOP3Bccs (Figures 5F and G). In parallel, ectopic expression of FLAG-tagged TRIM41 increased ubiquitylated TOP3Bccs (Figure 5H). Combined, these results show that TRIM41 plays a major role in the ubiquitylation and repair of TOP3Bccs.

### TRIM41-mediated ubiquitination and proteasomal degradation of TOP3Bccs promotes processing of TOP3Bccs by TDP2

Finally, we investigated the functional connection between TDP2-mediated processing of TOP3Bccs and the ubiquitin-proteasome pathway described above. Wild-type and TDP2KO HCT116 were transfected with FLAG-tagged R338W TOP3B alone or co-transfected with HA-tagged TDP2 and before harvesting were treated with the proteasome inhibitor MG132. RADAR assays demonstrated that while TDP2KO cells or treatment with MG132 increased cellular TOP3Bccs, proteasomal inhibition led to a greater accumulation of TOP3Bccs compared to TDP2 inactivation alone (Figures 6A-B). Combining these two conditions in cells ectopically expressing R338W TOP3B, i.e. blocking the proteasomal pathway and inactivating TDP2 (TDP2KO cells), produced no further increase in cellular TOP3Bcc levels compared to only MG132 treatment. Moreover, ectopic expression of HA-tagged TDP2 could not alter cellular TOP3Bcc levels in MG132-treated TDP2KO cells (Figures 6A-B). Together these results suggest that TDP2 and proteasomal degradation act in the same pathway for the excision of TOP3Bccs, with the ubiquitin-proteasome system working upstream to TDP2.

**Figure 6.**
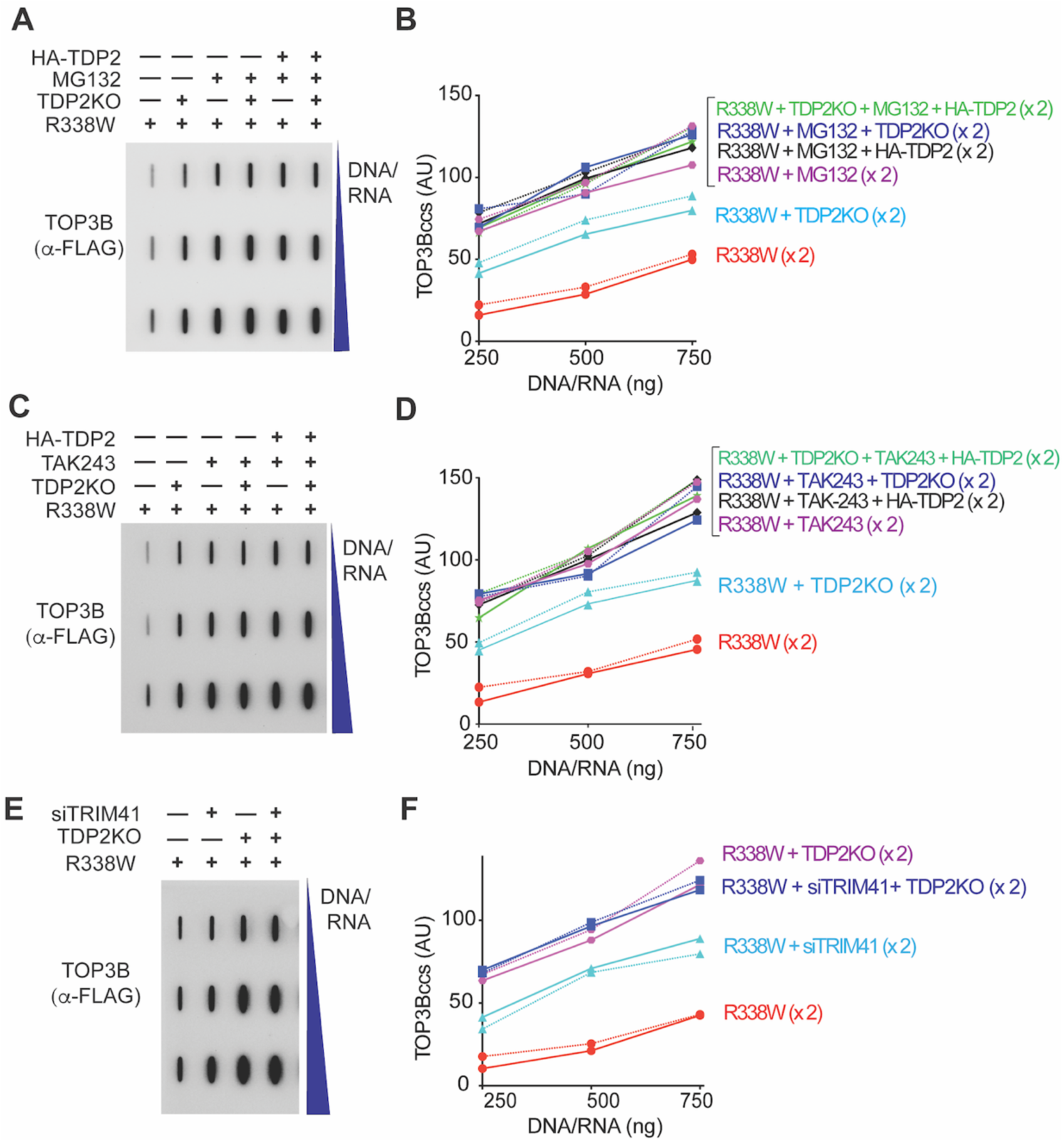
TDP2-Mediated Repair of TOP3Bccs is Dependent on Ubiquitination and Proteasomal Degradation. **(A)** Wild-type and TDP2KO HCT116 cells were transfected with FLAG-tagged R338W TOP3B alone or co-transfected with HA-tagged TDP2 plasmid construct and incubated for 72 h. Before harvest, cells were treated with MG132 (10 µM, 2 h). Nucleic acids and protein-nucleic acid adducts were recovered by RADAR assay, slot-blotted and TOP3Bccs were detected using anti-FLAG antibody. The figure is representative of two independent experiments. **(B)** Quantitation of TOP3Bccs in 2 independent RADAR assays as shown in panel A. TOP3Bccs were measured by densitometric analyses of slot-blot signals and plotted individually (x2) as a function of total nucleic acid (DNA and RNA) concentration. **(C)** Wild-type and TDP2KO HCT116 cells were transfected with FLAG-tagged R338W TOP3B alone or co-transfected with HA-tagged TDP2 plasmid construct and incubated for 72 h. Before harvest, cells were treated with TAK-243 (10 µM, 2 h). Nucleic acids and protein-nucleic acid adducts were recovered by RADAR assay, slot-blotted and TOP3Bccs were detected using anti-FLAG antibody. The figure is representative of two independent experiments. **(D)** Quantitation of TOP3Bccs in 2 independent RADAR assays as shown in panel C. **(E)** Wild-type and TDP2KO HCT116 cells were transfected with FLAG-tagged R338W TOP3B alone or co-transfected with siTRIM41 constructs and incubated for 72 h. Cells were harvested, nucleic acids containing protein adducts isolated by RADAR assay and slot blotted. TOP3Bccs were detected with anti-FLAG antibody. The figure is representative of two independent experiments. **(F)** Quantitation of TOP3Bccs in 2 independent RADAR assays as shown in panel E.

Next we performed experiments using the E1 ubiquitin-activating enzyme inhibitor (UAE1) TAK-243 (Hyer et al., 2018) instead of MG132. RADAR assays demonstrated that inhibition of ubiquitin conjugation by TAK-243 enhanced TOP3Bccs (compared to control and TDP2KO condition). Neither total depletion (knockout) nor overexpression of TDP2 could change TOP3Bcc levels in TAK-243-treated cells (Figures 6C-D). These results further indicate that, TOP3Bccs are ubiquitylated and proteolyzed by the proteasome before TDP2 could act.

Finally, we tested whether TRIM41, which we identified as E3 ligase for TOP3Bccs (see above), affects TDP2-mediated processing of TOP3Bccs. Wild-type and TDP2KO HCT116 cells were transfected with FLAG-tagged R338W TOP3B alone or co-transfected with siTRIM41 constructs and probed by RADAR assays. Compared to siTRIM41-treated cells, TDP2KO cells showed enhanced accumulation of TOP3Bccs. In addition TRIM41 depletion in TDP2KO cells could not further increase cellular TOP3Bcc levels (Figures 6E-F). Taken together, these results lead us to the conclusion that TRIM41 and TDP2 works in the same pathway for the processing of TOP3Bccs (Figure 7).

**Figure 7:**
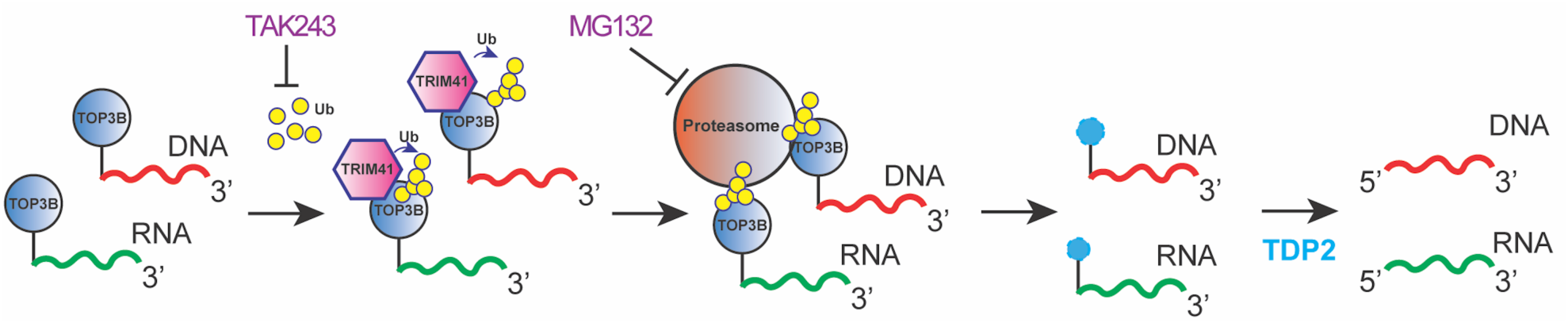
Schematic Representation of the Sequential repair of TOP3Bccs. Substitution of arginine 338 with tryptophan traps TOP3B both in DNA and RNA (TOP3Bccs) inside cells. Repair of trapped TOP3Bccs begins with ubiquitylation of TOP3B by TRIM41, followed by proteasomal degradation. TAK-243 inhibits ubiquitylation of TOP3Bccs and proteasomal degradation is blocked by MG132. Finally, TDP2 excises TOP3B polypeptide remaining attached to the 5’-end of DNA and RNA molecules.

## DISCUSSION

An outstanding question is whether RNA acquires abnormal topological structures during cellular metabolic processes requiring the activity of a topoisomerase. To our knowledge, our study provides the first direct *in vivo* evidence that TOP3B works enzymatically on cellular RNA. We demonstrate the isolation of catalytic intermediates of TOP3B and show that TOP3B can cleave both DNA and RNA by attaching itself covalently to the end of the cleaved nucleic acid to form DNA and RNA cleavage complexes (RNA and DNA TOP3Bccs). We also reveal the first pathway for the repair of stabilized TOP3Bccs in human cells. Both tyrosine phosphodiesterase 2 (TDP2) and the ubiquitin-proteasome system, with TRIM41 acting as E3 ubiquitin ligase for TOP3Bccs, come together to excise potentially lethal TOP3Bccs. Ubiquitination and proteasomal digestion of TOP3Bccs enable their processing by TDP2 (Figure 7).

The catalytic cycle of type IA topoisomerases consists in self-reversible TOPccs with the sequential: (i) cleavage of a single-stranded nucleic acid by formation of a covalent phosphodiester bond between the enzyme catalytic tyrosine and the 5’-end of the cleaved nucleic acid; (ii) resolution of the topological hindrance by passage of an intact strand through the topoisomerase-linked nucleic acid gate; and (iii) nucleic acid rejoining with release and turnover of the enzyme (Mills et al., 2018). To capture topoisomerase reaction intermediates (TOPccs), small molecules are used as molecular probes and therapeutic agents (antitumor and antibacterial). These small molecules (also called “topoisomerase poisons”) include camptothecin and its clinical derivatives topotecan and irinotecan for type IB topoisomerase [recently reviewed by (Thomas and Pommier, 2019)] and fluoroquinolones, doxorubicin, mitoxantrone and etoposide for type II topoisomerases (Maxwell, 1999; Nitiss, 2009).

For type IA topoisomerases there are no known small molecule inhibitors to act as “topoisomerase poisons”, which until now has made it impossible to detect catalytic intermediates of TOP3B trapped on DNA or RNA. However, screening studies with *Yersinia pestis* for SOS-inducing topoisomerase I mutations have identified point mutations in the TOPRIM motif and active site pocket of the enzyme that increase cellular levels of TOPccs (Cheng et al., 2009; Cheng et al., 2005; Narula et al., 2011). Corresponding mutations in *E. coli* topoisomerase I (Asp111Asn, Asp113Asn, Gly116Ser, Met320Val and Arg321Trp) also hamper cleavage-religation equilibrium of the enzyme and trap TOPccs. Multiple sequence alignment (E. coli Topo I, *Y. pestis* Topo I and Human TOP3B), showed that four of these residues are conserved in human TOP3B (Asp117, Asp119, Gly122 and Arg338). Mutating Arg338 to tryptophan and transient expression of the mutant construct in human cells resulted in significant *in vivo* accumulation of TOP3Bccs. Biochemical studies with the corresponding Arg321Trp mutant of *E. coli* topoisomerase I explained its altered behavior primarily as a defect in DNA rejoining of the TOPccs as well as a partial defect in DNA cleavage (Narula et al., 2011). Indeed, the positively charged conserved arginine residue, which is adjacent to the active site tyrosine and divalent magnesium ion is critical for the proper alignment and nucleophilic attack of the topoisomerase phosphotyrosyl bond by the 3’-hydroxyl-end of the cleaved nucleic acid (Narula et al., 2011). Our study shows that the corresponding arginine of TOP3B is also critical for the strand rejoining of TOP3Bccs both for DNA and RNA, and that substitution to tryptophan (R338W) produces a potent “self-trapping” topoisomerase.

Eukaryotic cells harbor two different tyrosyl DNA-phosphodiesterases, TDP1 and TDP2. Even though both hydrolyze topoisomerase tyrosyl-DNA phosphodiester bonds, they show no sequence homology or structural similarity (Bahmed et al., 2010; Cortes Ledesma et al., 2009; Kawale and Povirk, 2018; Nitiss and Nitiss, 2013; Nitiss et al., 2012; Pommier et al., 2014). By contrast to TDP1 (Cortes Ledesma et al., 2009; Yang et al., 1996), which belongs to the phospholipase D family (Interthal et al., 2001)], TDP2 belongs to the Mg^2+^/Mn^2+^-dependent family of endonucleases and exhibits high sequence similarities with the human DNA repair enzyme apurinic/apyrimidinic endonuclease-1 (APE1) (Cortes Ledesma et al., 2009; Gao et al., 2012; Nitiss and Nitiss, 2013; Pommier et al., 2014; Rodrigues-Lima et al., 2001). The 5’-tyrosyl-DNA phosphodiesterase activity of TDP2 makes TDP2 pivotal for the repair of trapped TOP2ccs in higher eukaryotes, and depletion of TDP2 increases cellular sensitivity to TOP2 “poisons” like etoposide (Cortes Ledesma et al., 2009; Gomez-Herreros et al., 2013; Maede et al., 2014; Zeng et al., 2011; Zeng et al., 2012). In addition, deleterious mutations of TDP2 are causative for spinocerebellar ataxia autosomal recessive 23 (SCAR23) (Zagnoli-Vieira et al., 2018).

Several *in vitro* and *in vivo* properties displayed by TDP2 made us hypothesize that it might resolve TOP3Bccs trapped in both DNA and RNA. Prior *in vitro* studies with purified human TDP2 indicated that, apart from processing double-stranded 5’-tyrosyl overhang substrate (mimicking TOP2ccs), TDP2 is most active with single-stranded DNA substrates bearing a 5′-phosphotyrosine terminus (mimicking DNA-TOP3Bccs) (Gao et al., 2012). These findings corroborate the crystal structures for TDP2 with its narrow nucleic acid groove optimal for binding single-stranded DNA ends (Schellenberg et al., 2012; Shi et al., 2012). Recombinant human TDP2 also has the ability to process a 5′-tyrosine covalently linked to a ribonucleotide or a polyribonucleotide (mimicking RNA-TOP3Bccs), and co-crystal structure of TDP2 bound to a tyrosyl-RNA substrate further confirms this finding (Gao et al., 2014). Finally, TDP2 is known to act as the VPg unlinkase that hydrolyzes the covalent phosphotyrosyl bond between the viral VPg protein and the 5′-end of viral RNAs (mimics of RNA-TOP3Bccs) during the replication cycle of viruses of the picornaviridae family (Kawale and Povirk, 2018; Pommier et al., 2014; Virgen-Slane et al., 2012). Biochemical and cellular assays performed in the present study reveal that DNA and RNA TOP3Bccs are both processed by TDP2 in human cells. But unlike the VPg protein, TDP2 can not excise TOP3Bccs unless TOP3B is denatured or proteolyzed. This finding is in line with the fact that purified TDP2 is also unable to process native TOP2ccs *in vitro* unless TOP2B is proteolyzed or denatured (Gao et al., 2014; Schellenberg et al., 2017).

Based on the present study, we propose that the proteasome-ubiquitin pathway enables TDP2 to access the TOP3B-DNA/ RNA phosphodiester bond of persistent TOP3Bccs. The 26S proteasome, a 2.5-MDa complex containing 33 subunits, is the major cellular proteolytic machine in the nucleus and cytosol (Bhattacharyya et al., 2014; Desai, 2012; Finley et al., 2016; Palmer et al., 1994; Rivett, 1998; Saeki, 2017). Ubiquitin, a small (76 amino acid) highly conserved globular protein, forms an isopeptide bond with a lysine residue of the substrate proteins that are destined for degradation by 26S proteasome (Desai, 2012; Hatakeyama, 2017; Komander and Rape, 2012; Saeki, 2017). The E1 ubiquitin-activating enzymes (UAE), E2 ubiquitin-transferring enzymes and E3 ubiquitin ligases act in concert to transfer mono- or poly-ubiquitin to substrate proteins (Saeki, 2017) with the E3 ubiquitin ligases playing a key role in substrate recognition (Hatakeyama, 2017). Ubiquitin forms polyubiquitin chains through one or more of its seven internal lysines (K6, K11, K27, K29, K33, K48 and K63) (Hatakeyama, 2017) with K48-linked polyubiquitin chains mainly targeting substrate proteins for proteasomal degradation (Chau et al., 1989; Johnson et al., 1995) (present study). Apart from canonical conjugates (K48-linked chains), polyubiquitin chains formed by K6, K11, K27, K29 or K63 can also promote proteasomal targeting and degradation (Bedford et al., 2011; Dammer et al., 2011; Jin et al., 2008; Kravtsova-Ivantsiv and Ciechanover, 2012; Meyer and Rape, 2014; Saeki et al., 2009; Xu et al., 2009).

Our results establish the importance of the proteasomal pathway for the repair of TOP3Bccs in addition to its role for both TOP1ccs and TOP2ccs in eukaryotic cells (Desai, 2012; Desai et al., 1997; Desai et al., 2003; Mao et al., 2001; Sordet et al., 2008; Zhang et al., 2006). We demonstrate that TOP3B is polyubiquitinated (involving K11, K27, K48 and K63 linkages) upon trapping on nucleic acids and processed by the 26S proteasome. Indeed, the UAE1 inhibitor TAK-243 and the proteasome inhibitor MG132 individually increase the cellular levels of TOP3Bccs induced by the R338W TOP3B self-poisoning mutant, consistent with the conclusion that the ubiquitin-proteasome pathway plays a critical role in removing aberrant TOP3Bccs. Further, inhibition of ubiquitin conjugation by TAK-243 or inhibition of the proteasome by MG132 trapped more TOP3Bccs than knocking-out TDP2KO, and overexpression of TDP2 could not change TOP3Bcc levels in TAK-243/ MG132 treated cells. These finding suggests that the ubiquitin-proteasome system and TDP2 act in a concerted way and proteolysis of trapped TOP3Bccs precedes their processing by TDP2.

We also demonstrate that TRIM41 (also known as RINCK) acts as E3 ubiquitin ligase for TOP3Bccs. TRIM41 is a member of the tripartite motif (TRIM) protein family, which includes more than 80 members in humans and is one of the largest families of the single-protein RING-type E3 ubiquitin ligases (Hatakeyama, 2017; Meroni and Diez-Roux, 2005; Napolitano and Meroni, 2012; Tanaka et al., 2005). Mounting evidence points to the fact that TRIM41 is a bona fide E3 ubiquitin ligase mediating ubiquitination and degradation of different substrates including protein kinase C (Chen et al., 2007) and the transcription factor ZSCAN21 (Lassot et al., 2018). In addition, TRIM41 plays an important role in antiviral innate immune responses by mediating the polyubiquitination and degradation of the influenza A virus nucleoprotein (Patil et al., 2018), the monoubiquitination of cGAS (Liu et al., 2018) and polyubiquitylation to limit hepatitis B virus replication (Zhang et al., 2013). Identification of TRIM41 as binding partner of TOP3B was previously reported (Kobayashi and Hanai, 2001; Rolland et al., 2014) but had not been connected to its role as E3 ubiquitin ligase. Experimental evidence provided by the current study points towards the fact that TOP3Bccs are ubiquitylated by TRIM41. Knocking-down TRIM41 reduced the ubiquitylation of TOP3Bccs, increased the stability of total cellular TOP3B and led to an accumulation of TOP3Bccs both in DNA and RNA. Conversely, overexpression of TRIM41 increased ubiquitylated TOP3Bccs and reduced cellular TOP3Bcc levels. Our study also reveals that TRIM41-mediated ubiquitylation and proteasomal degradation of nucleic acid-bound TOP3B further promotes processing of TOP3Bccs by TDP2.

In summary, our study provides the first direct evidence for TOP3B catalytically working (forming TOPccs) both with cellular RNA and DNA. It shows that TRIM41, the ubiquitin-proteasome system and TDP2 belong to a common repair pathway eliminating abnormal TOP3Bccs inside cells, and it set the stage for isolating and mapping TOP3Bccs in cellular DNA and RNA using the self-poisoning R338W TOP3B mutant. Our model for the sequential repair of TOP3Bccs is illustrated in Figure 7. Ubiquitylation of TOP3Bccs by TRIM41 leads to their degradation by the proteasome, which provides access for TDP2 to hydrolyzes the phosphotyrosyl bond and excise the TOP3B polypeptide remaining attached to the 5’-end DNA and RNA.

## ACKNOWLEGMENTS

We thank Protein Expression Laboratory [Protein and Nucleic Acid Production - Center for Cancer Research (CCR)], NCI-Frederick, MD for helping in production of recombinant human TOP3B. Our studies are supported by the Center for Cancer Research, the Intramural Program of the National Cancer Institute, NIH, Bethesda, Maryland 20892 (Z01 BC Z01 BC 006161-17 & Z01 BC 006150-19).

## AUTHOR CONTRIBUTIONS

Y.P. supervised the study. S.S. and Y.P. devised the concept. Y.P., S.S., Y.S., S.-Y.H. and H.Z. designed the experiments. S.S., Y.S., S.-Y.H. and U.J. performed experiments and data analysis. S.S. and Y.P. wrote the manuscript.

## DECLARATION OF INTERESTS

The authors declare no competing interests.

## STAR⋆METHODS

### KEY RESOURCES TABLE

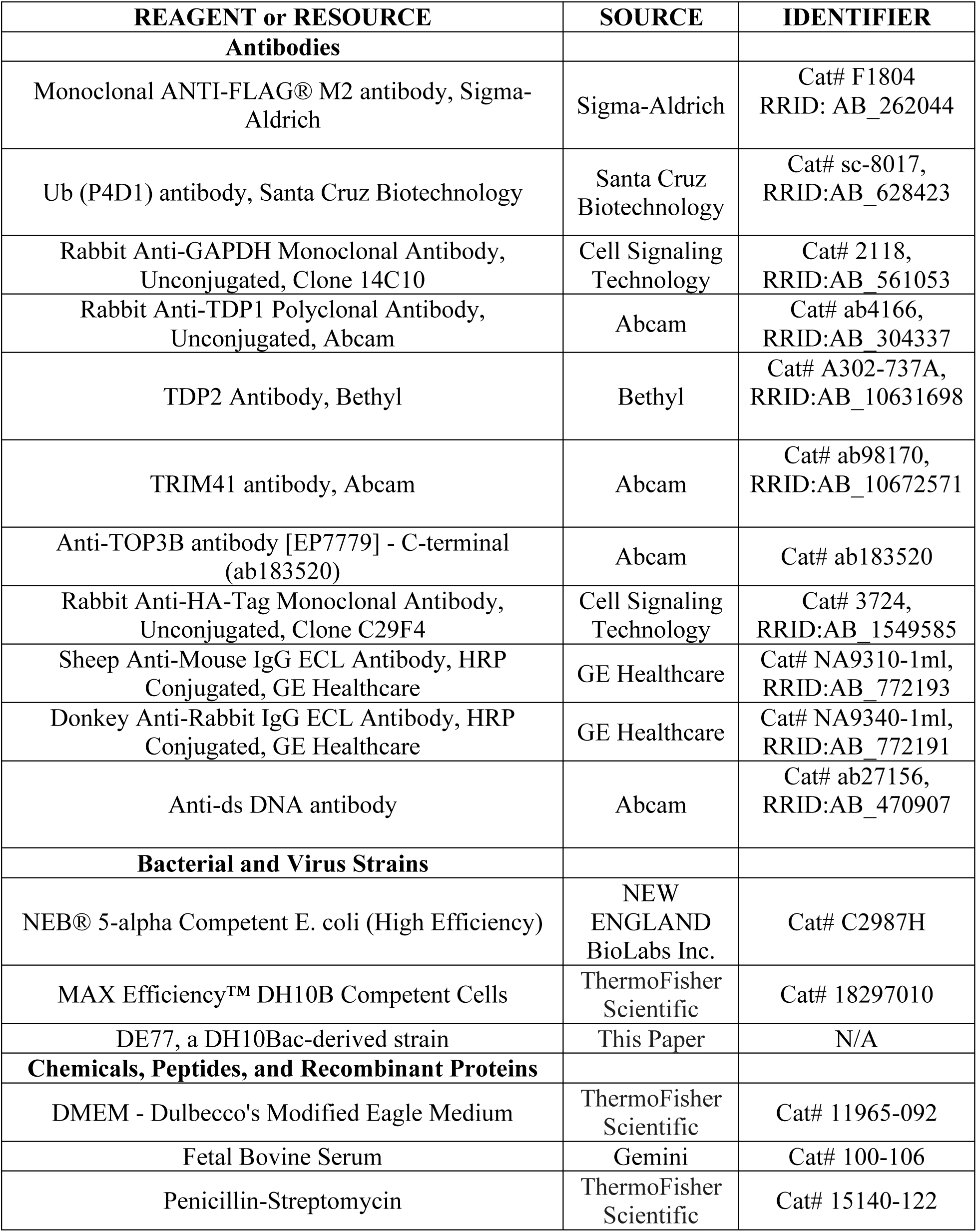
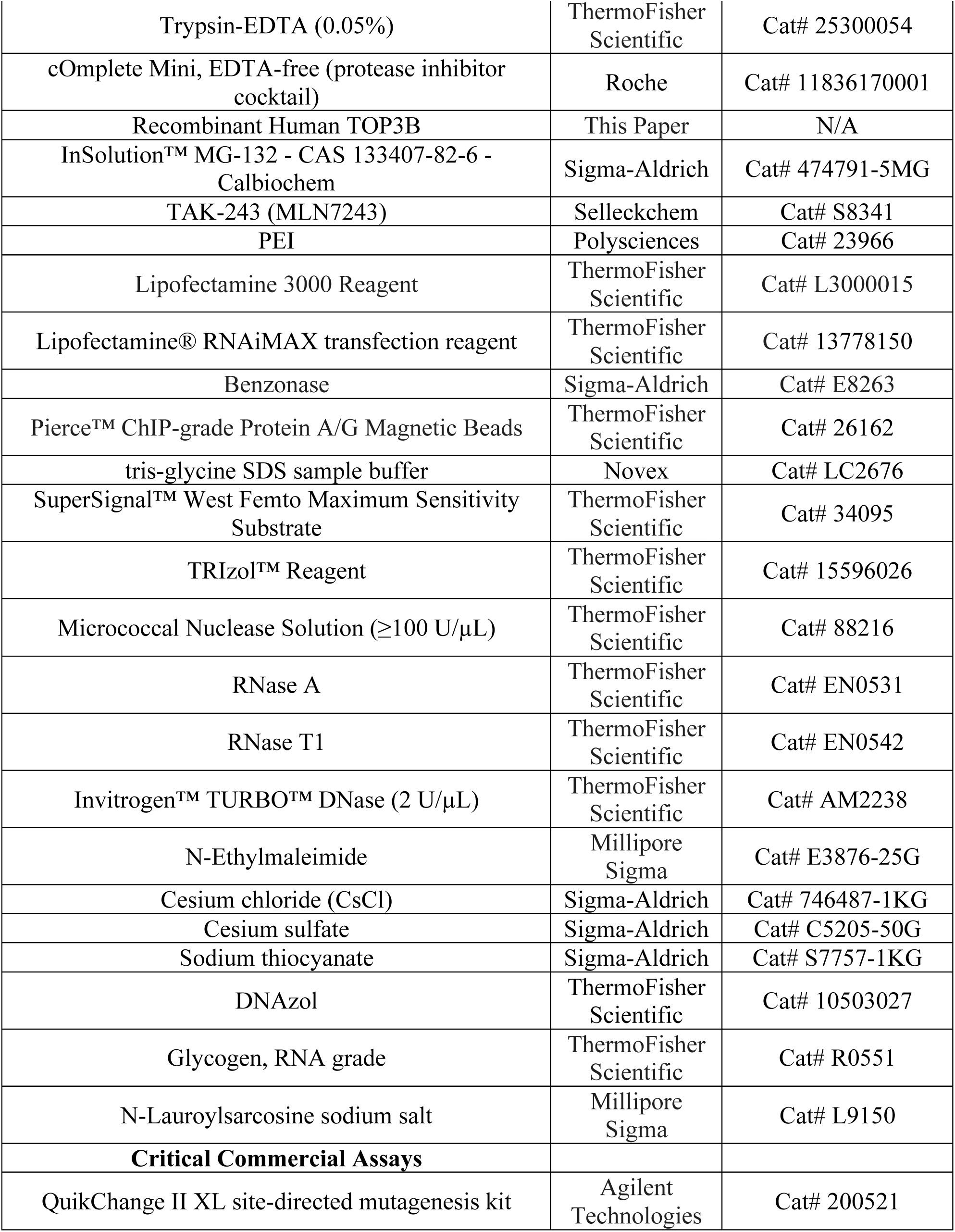
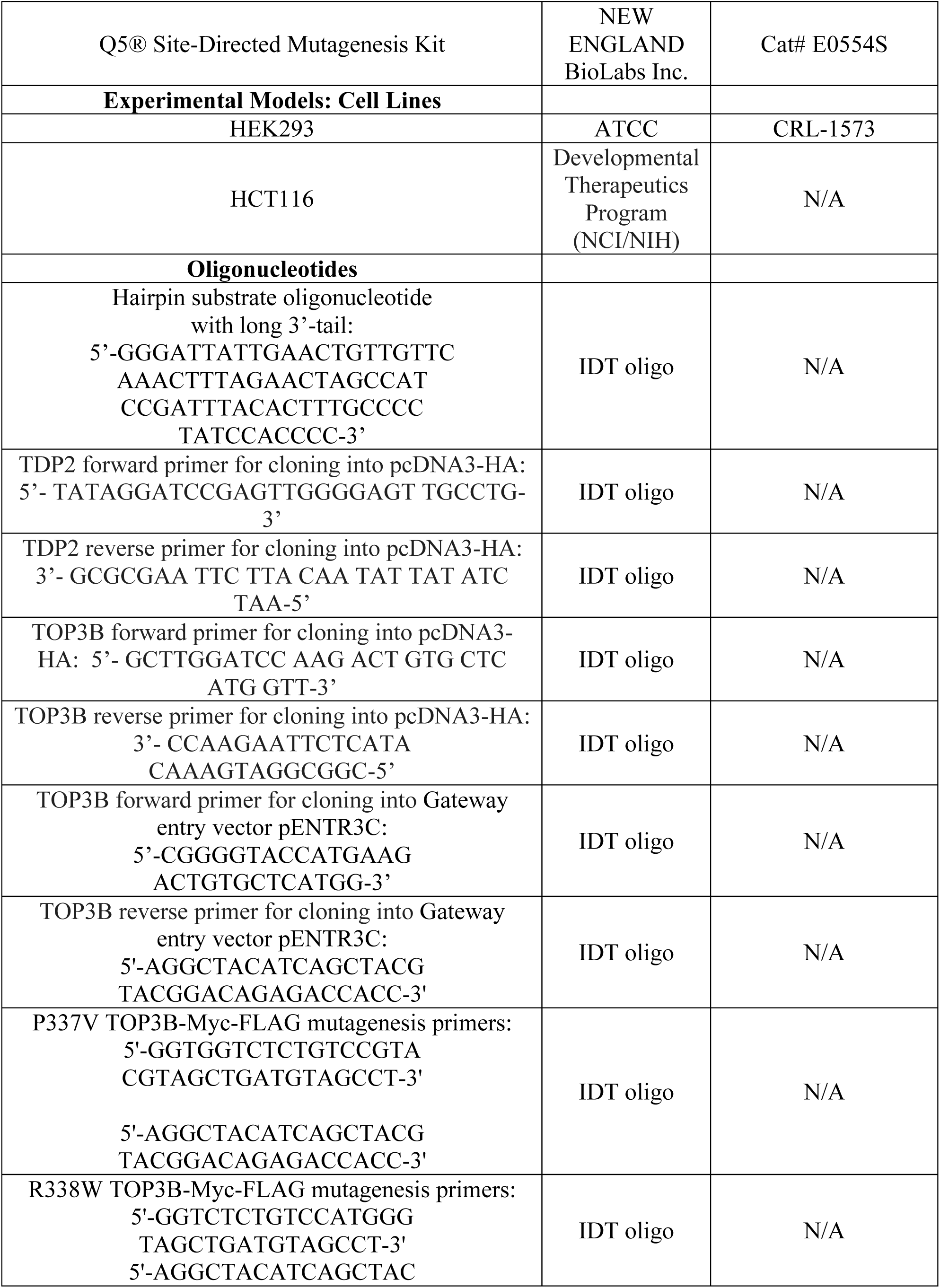
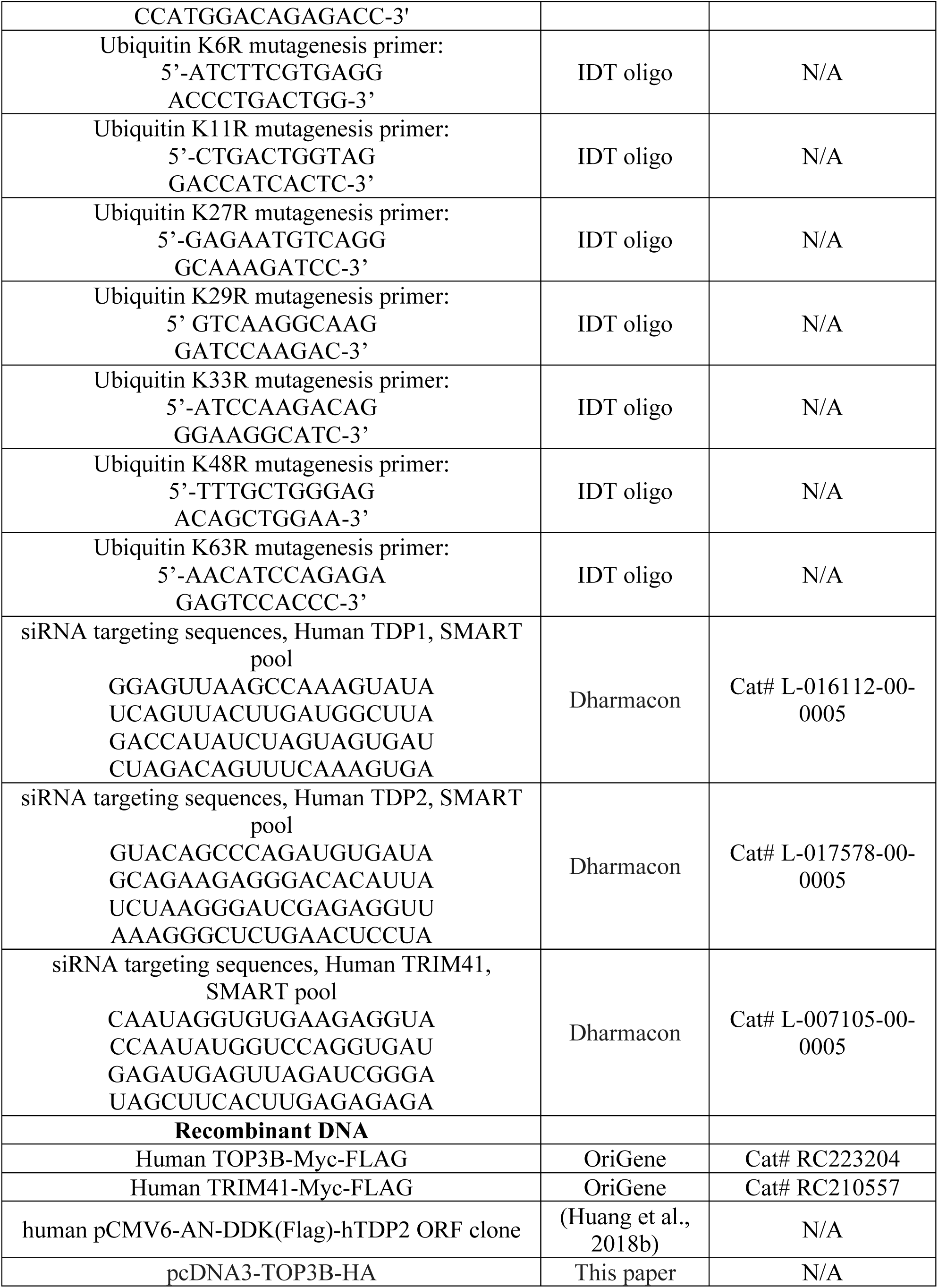
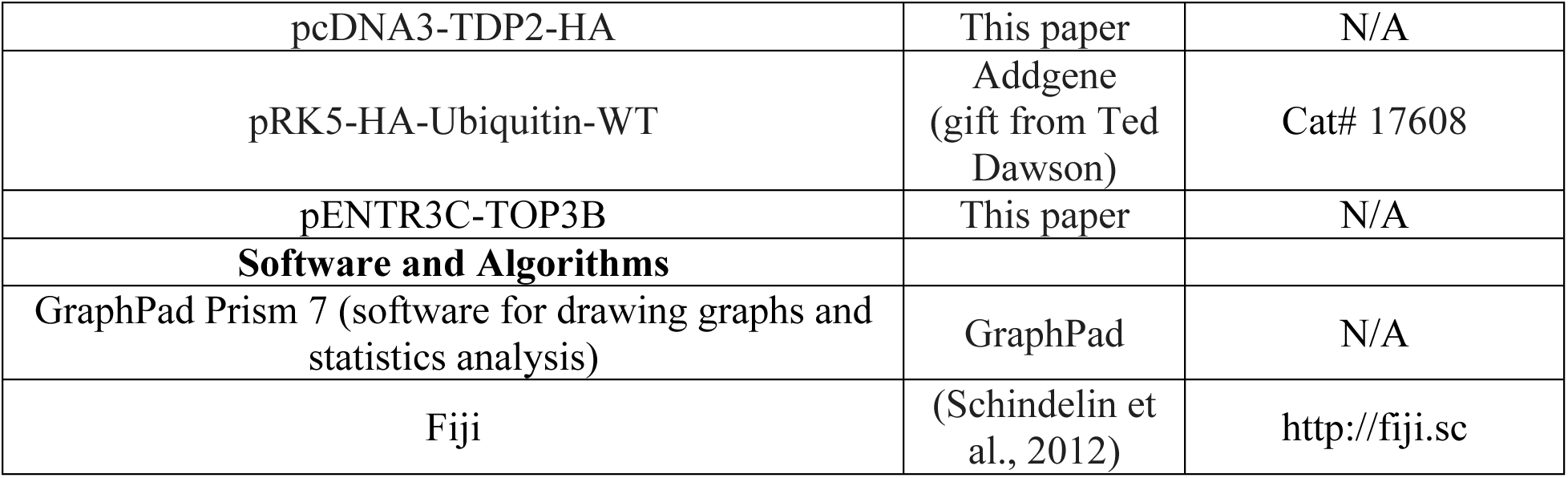

## CONTACT FOR REAGNET AND RESOURCE SHARING

Further information and requests for reagents may be directed to and will be fulfilled by Lead Contact Yves Pommier.

## METHODS DETAILS

### Cell culture

HEK293 (ATCC, Manassas, VA) and HCT116 (Developmental Therapeutics Program, National Cancer Institute) cell lines were grown in Dulbecco’s modified Eagle’s medium (Life Technologies, Carlsbad, CA) supplemented with 10% Fetal Bovine Serum (Gemini, West Sacramento, CA) and 1% penicillin-streptomycin (ThermoFisher Scientific, 15140122) at 37°C in humidified 5% CO2 chamber. HCT116 TDP2 knockout (TDP2KO) cells were generated by CRISPR/Cas9 method as previously described (Huang et al., 2018b).

### Mammalian Expression Constructs and Transient Expression in Mammalian Cell

Human TOP3B-Myc-FLAG cDNA ORF (CAT#: RC223204) and Human TRIM41-Myc-FLAG cDNA ORF (CAT#: RC210557) Clones were purchased from OriGene. The full-length cDNAs of TDP2 and TOP3B were PCR-amplified from human pCMV6-AN-DDK(Flag)-hTDP2 ORF clone (Huang et al., 2018b) and TOP3B-Myc-FLAG cDNA ORF Clone (CAT#: RC223204) respectively using cloning primers (TDP2 forward primer 5’-TATAGGATCCGAGTTGGGGAGTTGCCTG-3’; TDP2 reverse primer 3’-GCGCGAATTCTTACAATATTATATCTAA-5’; TOP3B forward primer 5’-GCTTGGATCCAAGACTGTGCTCATGGTT-3’; TOP3B reverse primer 3’-CCAAGAATTC TCATACAAAGTAGGCGGC-5’) and subcloned into pcDNA3-HA with BamHI and EcoRI sites. HA-Ubiquitin WT plasmid was a gift from Ted Dawson (Addgene plasmid CAT#: 17608). Plasmids were transfected in HCT116 and HEK293 cells using Lipofectamine 3000 Reagent (CAT#: L3000015, ThermoFisher Scientific) according to the manufacturer’s protocol for 48–72 h.

### Recombinant Human TOP3B Production

TOP3B was initially PCR amplified from Human TOP3B-Myc-FLAG cDNA ORF (CAT#: RC223204) using forward primer: 5’-CGGGGTACCATGAAGACTGTGCTCATGG-3’ and reverse primer: 5’-CCGCTCGAGTCATACAAAGTAGGCGGCCAG-3’ and cloned into Gateway entry vector pENTR3C (Invitrogen, CAT#: A10464). TOP3B was then subcloned by Gateway LR recombination (Thermo Fisher) into pDest-635 (22876-X01-635) for insect cell expression which includes an N-terminal His6 tag. Bacmid was prepared in DE77, a DH10Bac-derived strain (Bac-to-Bac system, Thermo Fisher) and after purification, bacmid DNA was verified by PCR amplification across the bacmid junctions. Bacmids were transfected in SF-9 cells using PEI (1 mg/ml with 5% glucose; Polysciences, CAT#: 23966), recombinant baculovirus stock was collected and titrated using ViroCyt (Beckamn). Two liters of Tni-FNL cells were set in a baffled 5-liter Thomson Optimum Growth Flask in Gibco Express 5 medium with 18mM glucose at a cell density of 1 × 10^6^ cells/ml at 27°C and 24 hrs later infected at a MOI (multiplicity of infection) of 3. After 3 days of incubation at 21°C, cell pellets were collected by centrifugation at 2000 rpm for 11 min and flash frozen on dry ice. Cell pellet was thawed by the addition of 200 ml of lysis buffer (20 mM HEPES, 300 mM NaCl, 1 mM TCEP and 1:100 v/v of Sigma protease inhibitor P8849) and homogenized by vortexing. The cells were lysed by performing two passes on an M-110EH-30 microfluidizer (Microfluidics) at 7000 psi, clarified at 100K x g for 30 minutes at 4°C using an optima L-90K ultracentrifuge (Beckman), filtered (0.45 micron) and applied to a f20 mL IMAC HP column (GE Scientific) that was pre-equilibrated with lysis buffer containing 50 mM imidazole on a Bio-Rad NGC. Column was washed with lysis buffer containing 50 mM imidazole and proteins were eluted with lysis buffer containing 500 mM imidazole. After SDS-PAGE/Coomassie staining, positive fractions were pooled, dialyzed to 20 mM HEPES, 50 mM NaCl, 1 mM TCEP, 0.5 mM PMSF, 1:1000 v/v of PI, 50% glycerol, pH 7.2. Protein concentration was determined (0.88 mg/ml) and stored at -80°C for future use.

### siRNA Transfection

Silencing of TDP1, TDP2 and TRIM41 were done using ON-TARGETplus SMARTpool siRNA targeting TDP1 (CAT#: L-016112-00-0005), TDP2 (CAT#: L-017578-00-0005) and TRIM41 (CAT#: L-007105-00-0005) respectively. All siRNAs were used at a final concentration of 25 nM and transfected using Lipofectamine® RNAiMAX transfection reagent (CAT#: 13778150, ThermoFisher Scientific) following the manufacturer’s protocol for 48–72 h.

### Site-Directed Mutagenesis (SDM) in Mammalian Expression Vectors

Site-Directed Mutagenesis was performed using QuikChange II XL site-directed mutagenesis kit (Agilent Technologies) following the manufacturer’s protocol. P337V TOP3B-Myc-FLAG was generated using oligonucleotides: 5’-GGTGGTCTCTGTCCGTACGTAGCTGATGTAGCCT-3’ and 5’-AGGCTACATCAGCTACGTACGGACAGAGACCACC-3’. R338W TOP3B-Myc-FLAG was generated using oligonucleotides: 5’-GGTCTCTGTCCATGGGTAGCTGATGTAGCCT-3’ and 5’-AGGCTACATCAGCTACCCATGGACAGAGACC-3’. R338W TOP3B-HA was generated using oligonucleotides: 5’-GGTCTCTGTCCATGGGTAGCTGATGTAGCCT-3’ and 5’-AGGCTACATCAGCTACCCATGGACAGAGACC-3’. Ubiquitin K6R was generated by Q5 SDM Kit using oligonucleotide 5’-ATCTTCGTGAGGACCCTGACTGG-3’. Ubiquitin K11R was generated using oligonucleotide 5’-CTGACTGGTAGGACCATCACTC-3’. Ubiquitin K27R was generated using oligonucleotide 5’-GAGAATGTCAGGGCAAAGATCC-3’. Ubiquitin K29R was generated using oligonucleotide 5’ GTCAAGGCAAGGATCCAAGAC-3’. Ubiquitin K33R was generated using oligonucleotide 5’-ATCCAAGACAGGGAAGGCATC-3’. Ubiquitin K48R was generated using oligonucleotide 5’-TTTGCTGGGAGACAGCTGGAA-3’. Ubiquitin K63R was generated using oligonucleotide 5’-AACATCCAGAGAGAGTCCACCC-3’.

### Western Blotting and antibodies

To prepare whole cell lysates for Western blotting, cells were resuspended with RIPA buffer (150 mM NaCl, 1% NP-40, 0.5% Sodium deoxycholate, 0.1% SDS, 50 mM Tris pH 7.5, 1 mM DTT and protease inhibitor cocktail). After thorough mixing, samples were agitated at 4°C for 30 min, sonicated for 30 seconds with 50% pulse, centrifuged at 15,000 × g at 4°C for 15 min, and supernatants were collected.

Lysed samples were mixed with tris-glycine SDS sample buffer (Novex, LC2676) and loaded onto Novex tris-glycine gels (Novex). Blotted membranes were blocked with 5% non-fat dry milk in PBS with 0.1% Tween-20 (PBST). Primary antibodies were diluted in 5% milk in PBST by 1:1000 for Mouse monoclonal anti-FLAG M2 (Sigma-Aldrich, St. Louis, MO, CAT#: F1804), Mouse monoclonal anti-Ub (P4D1) antibody (Santa Cruz Biotechnology, Dallas, Texas, CAT#: sc-8017), 1:10000 for Rabbit anti-GAPDH monoclonal antibody (Cell Signaling Technology, Danvers, MA, CAT#: 2118S), 1: 500 for Rabbit polyclonal anti-TDP1 (Abcam, Cambridge, MA, CAT#: ab4166,), 1: 500 for Rabbit polyclonal anti-TDP2 (Bethyl, Montgomery, TX, CAT#: A302-737A), 1:500 for Rabbit polyclonal anti-TRIM41 (Abcam, Cambridge, MA, CAT#: ab98170), 1:1000 for Rabbit monoclonal anti-HA (Cell Signaling Technology, Danvers, MA, CAT#: 3724S). Secondary antibodies were diluted (1:10000) in 5% non-fat milk in PBST and signal was detected by ECL chemiluminescence reaction (Thermo Scientific, Waltham, MA).

### ICE bioassay

TOP3B-DNA and -RNA cleavage complexes were isolated using the in vivo complex of enzyme (ICE) bioassay (Pourquier et al., 2000; Shaw et al., 1975). Briefly, FLAG-tagged R338W TOP3B transfected cells were pelleted and immediately lysed with 1 ml of 1% sarkosyl. After homogenization with a Dounce, cell lysates were gently layered on step gradients containing four different CsCl (Sigma-Aldrich, CAT#:746487-1KG) solutions (2 ml of each) of the following densities: 1.82, 1.72, 1.50, and 1.45 (Shaw et al., 1975). The gradients were prepared by diltuting a stock solution of CsCl of density 1.88. Cesium sulfate (Sigma-Aldrich, CAT#:C5205-50G) was included in the bottom solution of density 1.82 to help in flotation of the RNA and sodium thiocyanate (Sigma-Aldrich, CAT#:S7757-1KG) was included in topmost solution of density 1.45 to facilitate the complete removal of non-covalently bound proteins from the sedimenting nucleic acid species (Shaw et al., 1975). Tubes were centrifuged at 30,700 rpm in a Beckman SW40 rotor for 24 h at 20°C. Half-milliliter fractions were collected from the bottom of the tubes. Fractions containing DNA and RNA were pooled separately, quantitated, diluted with 25 mM sodium phosphate buffer (pH 6.5), and applied to Immobilon-FL PVDF 0.45 μm membranes (Merck Millipore, USA, CAT#: IPFL00010) through a slot-blot vacuum manifold as described (Pourquier et al., 2000). TOP3Bccs were detected with the Mouse monoclonal anti-FLAG M2 antibody (Millipore Sigma, St. Louis, MO, CAT#: F1804).

### RADAR Assay

FLAG-tagged R338W TOP3B transfected cells (1 × 10^6^) were washed with PBS and lysed by adding 1 ml DNAzol (ThermoFisher Scientific, CAT#:10503027). Nucleic acids were precipitated following addition of 0.5 ml of 100% ethanol, incubation at -20°C for 5 min and centrifugation (12,000 × g for 10 min). Precipitates were washed twice in 75% ethanol, resuspended in 200 μl TE buffer, heated at 65°C for 15 minutes, followed by shearing with sonication (40% power for 15 sec pulse and 30 sec rest 5 times). Samples were centrifuged at 15,000 rpm for 5 min and the supernatant containing nucleic acids with covalently bound proteins were collected. Nucleic acid containing protein adducts were quantitated, slot-blotted and TOP3Bccs were detected with Mouse monoclonal anti-FLAG M2 antibody (Millipore Sigma, St. Louis, MO, CAT#: F1804).

### Detection of Ubiquitylated and SUMOylated TOPccs (DUST) Assay

For the DUST assay (Sun et al., 2019), nucleic acids and covalent protein-nucleic acid adducts were recovered from FLAG-tagged R338W TOP3B transfected cells using the RADAR assay. 8 μg of each RADAR assay sample was digested with 250 units micrococcal nuclease (Thermo Fisher Scientific, 100 units/μl) in the presence of 5 mM CaCl2, followed by SDS-PAGE electrophoresis for immunodetection of total TOP3Bccs and ubiquitylated TOP3Bccs. In addition, each RADAR sample was subjected to slot-blotting and immunodetection with anti-dsDNA antibody (Abcam, ab27156) to confirm equal DNA loading.

### Isolation of Cellular Covalent RNA-protein Adducts from Cells Using TRIzol® Reagent

RNA-protein adducts were isolated from FLAG-tagged R338W TOP3B transfected cells as described (Trendel et al., 2018). Briefly, 1 × 10^7^ cells were lysed in 1 ml TRIzol™ Reagent (Invitrogen, USA, CAT#:15596026) by pipetting the samples up and down several times followed by incubation at room temperature for 5 min. 200 μl chloroform was added to the samples and mixed properly by inverting the tubes. After incubation at room temperature for 3 min and centrifugation for 10 minutes at 7,000 × g at 4°C, the aqueous phase was removed, and the interphase was transferred to a new tube. The interphase was gently washed twice with 1 ml low SDS buffer (Tris-Cl 50 mM, EDTA 1 mM, SDS 0.1 %), resuspended in low SDS buffer, centrifuged at 5000 × g for 2 min at room temperature and the supernatant was stored. Pellets were dissolved again with 1 ml of low SDS buffer, then twice with 1 ml high SDS buffer (Tris-Cl 50 mM, EDTA 1 mM, SDS 0.5 %) and all the supernatants were stored following centrifugation. NaCl was added to a final concentration of 300 mM to each of the interphase eluates, along with 10 ug of RNase-free glycogen and 1 ml isopropanol before mixing by inversion. Samples were spun down for 15 min with 18,000 × g at -10 °C. Supernatant were discarded, pellet was resuspended in 70 % ethanol. Samples were again centrifuged for 1 min at 18000 × g at room temperature. Supernatant were discarded, residual ethanol removed, and the pellets were resuspended in nuclease-free water at 4°C. 10X TURBO DNase Buffer (ThermoFisher Scientific, USA) was added to the resuspended samples to 1X concentration along with 10 μl TURBO DNase (ThermoFisher Scientific, USA) and incubated for 60 minutes at 37°C and 700 rpm shaking. After DNase treatment, samples were isopropanol precipitated in the presence of 300 mM NaCl and dissolved in DEPC treated water. RNA purity and concentrations were estimated by spectroscopy on a NanoDrop 1000 Spectrophotometer (ThermoFisher Scientific, USA). Samples were slot-blotted and TOP3Bccs on RNA were detected using Mouse monoclonal anti-FLAG M2 antibody (Millipore Sigma, St. Louis, MO, CAT#: F1804).

### Generation of TOP3Bccs *in vitro* and digestion by TDP1, TDP2 and Benzonase

The hairpin substrate oligo nucleotide with long 3’-tail: GGGATTATTGAACTGTTGTTCAAACTTTAGAACTAGCCATCCGATTTACACTTTGCCCCTATCCACCCC was synthesized by IDT. 300 nM of annealed substrate was combined with 4 uM of purified recombinant TOP3B in 100 mM potassium glutamate (pH 7.0), 3 mM MgCl2, 0.02% v/v Tween-20, 1 mM DTT, and incubated at 30°C for 15 mins before addition of 1 or 3 μM of TDP1 or TDP2 and incubated for an additional 60 min at 25°C. Benzonase (3 or 9 Units) was used as positive control. SDS (0.2 %) was added to the samples to stop the reaction. The samples were resolved on 6% tris-glycine-SDS PAGE and Western blotting was carried out using standard techniques with rabbit monoclonal anti-TOP3B (Abcam, CAT#: ab183520).

## SUPPLEMENTAL FIGURE LEGENDS

**Supplemental Figure 1. TOP3Bccs form both in DNA and RNA**

**(A)** HEK293 cells were transfected with FLAG-tagged R338W TOP3B plasmid construct. TOP3Bccs in DNA and RNA are represented in red and green, respectively. Nucleic acids and protein-nucleic acid adducts were isolated by RADAR assay and equal amounts of RADAR assay samples were digested either with excess RNase A (200 μg/mL) and RNase T1 (200 Units/ml), with DNase I (10 Units/reaction) or with micrococcal nuclease (MNase, 300 Units/reaction). Samples were ethanol-precipitated, resuspended, slot-blotted and TOP3Bccs were detected using anti-FLAG antibody. The figure is representative of two independent experiments.

**(B)** Graphical representation of TOP3Bcc formation in 2 independent (x 2) RADAR assays as shown in panel A. TOP3Bccs were measured by densitometric analyses of slot-blot signals and plotted as a function of total nucleic acid (DNA and RNA) concentration.

**Supplemental Figure 2. TDP2 Processes both RNA and DNA TOP3Bccs**

**(A-B)** Western blots showing the efficiency of TDP1 and TDP2 knock-down (GAPDH as loading control). HEK293 cells were co-transfected with R338W TOP3B and siRNA constructs against TDP1, TDP2 or both TDP1 and TDP2, as indicated. Cell lysates were subjected to Western blotting with anti-TDP1 and anti-TDP2 antibody.

**(C)** Control Western blot showing no expression of TDP2 in TDP2KO HCT116 cells. Wild-type or TDP2KO HCT116 cells were transfected with FLAG-tagged R338W TOP3B and incubated for 72 h. Cell lysates were subjected to Western blotting and TDP2 was detected with anti-TDP2 antibody.

**(D)** HEK293 cells were transfected with R338W TOP3B construct alone or co-transfected with siTDP2 construct. After 72 h nucleic acid-containing protein adducts were isolated by RADAR assay and equal amounts of samples were digested either with excess RNase A (200 μg/mL) and RNase T1 (200 Units/ml), or with DNase I (10 Units/reaction). Samples were ethanol precipitated, resuspended, slot-blotted and TOP3Bccs were detected using anti-FLAG antibody. The figure is representative of two independent experiments.

**(E)** Quantitation of TOP3Bcc formation in 2 independent RADAR assays (x 2) as shown in panel D. TOP3Bccs were measured by densitometric analyses of slot blot signals and plotted as a function of total nucleic acid (DNA and RNA) concentration.

**(F)** Western blots showing expression levels of endogenous TDP2 and ectopically expressed HA-TDP2. Wild-type (WT) and TDP2KO HCT116 cells were transfected with FLAG-tagged R338W TOP3B alone or co-transfected with HA-tagged TDP2 plasmid construct and incubated for 72 hs. Cell lysates were subjected to Western blotting and immunoblotted using anti-TDP2 antibody.

